# PTPN14, a modifier of HHT, protects SMAD4 from ubiquitination and turnover to potentiate BMP9 signaling in endothelial cells

**DOI:** 10.1101/2021.09.29.462397

**Authors:** Ons Mamai, Daniah T. Beleford, Mark Taylor, Sugandha Basu, Xinjian Cen, Suprita Trilok, Jiamin Zhang, Allan Balmain, Rosemary J. Akhurst

**Affiliations:** Helen Diller Family Comprehensive Cancer Center, University of California, San Francisco, San Francisco, CA, USA; Department of Pediatrics, Division of Medical Genetics, University of California at San Francisco, San Francisco, CA, USA; Department of Biochemistry and Biophysics, University of California at San Francisco, San Francisco, CA, USA; Department of Cancer Genetics and Genomics, Roswell Park Comprehensive Cancer Center, Buffalo, NY, USA; Department of Anatomy, University of California at San Francisco, San Francisco, CA, USA

## Abstract

Hereditary Hemorrhagic Telangiectasia (HHT) results from germline loss-of-function mutations of *ENG, ACVRL1*, or *SMAD4*, encoding TGFβ/BMP signaling components. Telangiectasias occur in most patients, and pulmonary, visceral, or cerebral arteriovenous malformations (AVMs) in 20-50% of these. How HHT mutations cause these clinical manifestations and why some patients suffer more serious sequelae than others is unknown. *PTPN14* is a genetic modifier of pulmonary AVM incidence, and here we show by gene expression network analysis of a large panel of genetically diverse mouse lung RNA samples, that *Ptpn14* is ontologically associated with markers of angiogenesis, vascular remodeling, and BMP/TGFβ and Rho kinase signaling. We demonstrate physical interaction between protein tyrosine phosphatase non-receptor, type 14 (PTPN14) and SMAD4 in nucleus and cytoplasm of primary human endothelial cells. PTPN14 suppresses ubiquitination and turnover of SMAD4 to augment tonic SMAD-mediated transcriptional readouts. This is the first report that PTPN14 binds and stabilizes SMAD4, a key component of the HHT signaling pathway. Through this mechanism, and its inhibition of YAP/TAZ signaling, PTPN14 levels may protect against development of AVMs in HHT. We discuss potential druggable targets for HHT within the *ENG-ALK1-SMAD4-PTPN14* network.

**One Sentence Summary:** PTPN14 binds and stabilizes SMAD4 to potentiate BMP9 signaling in endothelial cells and components of the PTPN14 network may be drug targets for HHT.

## INTRODUCTION

Hereditary Hemorrhagic Telangiectasia (HHT) is an autosomal dominant genetic condition with an incidence of 1 case per 5,000 worldwide. More than 80% of cases are attributed to germline heterozygous loss of function (LOF) mutations in either the *ENG* (HHT type 1) or *ACVRL1* (HHT type 2) genes, which encode endoglin and activin receptor-like kinase-1 (ALK1), respectively (Faughnan *et al*, 2011; Shovlin, 2010). Heterozygous LOF mutations in *SMAD4* which also cause Juvenile Polyposis Syndrome (JPS/HHT) account for some 2% of HHT cases, and additional loci have been implicated in HHT including *GDF2*, that encodes BMP9 (Liu *et al*, 2020; Wooderchak-Donahue *et al*, 2013). Damaging genetic variants in *ENG, ACVRL1* and *SMAD4* have been added to the American College of Medical Genetics list of reportable secondary exome sequence findings, due to the likelihood of severe disease or death that may be preventable if identified before symptoms arise (Miller *et al*, 2021).

HHT patients frequently present during adolescence with mucocutaneous telangiectasias and frequent episodes of epistaxis (Faughnan *et al*., 2011; Shovlin, 2010). Telangiectasias arise through abnormal vascular remodeling, causing capillary breakdown and direct shunting from arterioles into venules without intervening capillary beds (Guttmacher *et al*, 1995). The blood vessel walls of telangiectasias are dilated and weak resulting in frequent hemorrhage and epistaxis. Bleeding from gastrointestinal tract telangiectasias can cause chronic anemia that requires repeated intravenous iron administration or blood transfusion. Approximately half of HHT patients also develop larger arteriovenous malformations (AVMs) in lung, brain, and liver that can have more serious clinical sequelae (Lesca *et al*, 2007; Letteboer *et al*, 2006). Direct shunting of blood from artery to vein in larger AVMs can cause local tissue hypoxia and ischemia, and AVMs are prone to hemorrhage. Brain AVMs are often asymptomatic but can cause sudden seizures or hemorrhage. Pulmonary AVMs may cause chronic hypoxemia and exercise intolerance due to impaired gas exchange, and stroke or brain abscess due to pulmonary right to left shunting that can cause paradoxical emboli of both thrombotic and septic origin.

The known HHT causative genes encode components of the TGFβ/BMP signal transduction pathway. Endoglin is a trans-membrane glycoprotein related to the TGFβ type III receptor, betaglycan. It binds several TGFβ superfamily ligands, including BMP9 and TGFβ1 (Barbara *et al*, 1999) and has two alternative spliced protein isoforms that differ in their short cytoplasmic tails (Velasco *et al*, 2008). The *ENG* gene is positively regulated by BMP/SMAD signaling (Calvo-Sánchez *et al*, 2019; Morikawa *et al*, 2011; Tual-Chalot *et al*, 2014). Under pathological situations, such as preeclampsia or inflammation, endoglin may be shed from the cell surface resulting in soluble circulating S-endoglin (Hawinkels *et al*, 2010; Venkatesha *et al*, 2006) that can act as a ligand trap. Endoglin regulates endothelial cell (EC) shape (Fernandez *et al*, 2005), directional migration of ECs against blood flow, and arteriovenous cell fate, consequently modulating EC location and maintenance of arterial identity (Jin *et al*, 2017; Sugden *et al*, 2017). ALK1 is a transmembrane activin type I receptor-like kinase that binds TGF and BMPs, with preferential binding and activation by BMP9 (David *et al*, 2007). After ligand binding, ALK1 phosphorylates the receptor-associated SMADs (R-SMADs) of the BMP pathway, SMAD1, 5 and 8. Finally SMAD4, the common co-SMAD that binds phosphorylated R-SMADs of both the BMP (SMAD1, 5 and 8) and TGFβ/activin/nodal (SMAD2 and 3) signaling pathways, forms a heterohexameric complex with activated R-SMAD1/5, which shuttles to the nucleus to instigate context-dependent transcriptional outputs (Derynck & Budi, 2019).

Although the HHT causative genes have been identified and characterized (Faughnan *et al*., 2011; Shovlin, 2010; Wooderchak-Donahue *et al*., 2013), knowledge of the molecular pathways leading to HHT lesions is still incomplete. Filling this knowledge gap might ultimately result in novel therapeutic strategies for HHT or other vascular disorders. Towards this goal, we previously searched for natural human genetic variants that protect or predispose to more serious HHT manifestations, such as AVMs, which occur in 10-50% of patients with HHT (Lesca *et al*., 2007; Letteboer *et al*., 2006). Screening 100 Dutch HHT families, we found genetic association between the presence of pulmonary AVMs and single nucleotide polymorphisms (SNPs) within the *PTPN14* gene (Benzinou *et al*, 2012). Based on this finding, we proposed that PTPN14 functionally interacts with the BMP9/ALK1 pathway, but the molecular mechanisms of this interaction have not yet been reported.

PTPN14 (protein-tyrosine phosphatase, non-receptor-type, 14, OMIM 613611) is a member of both the FERM family and the non-receptor tyrosine phosphatase family of proteins. Its carboxy-terminal phosphatase domain has been reported to dephosphorylate targets of Src kinase, including β-catenin, VE-cadherin, caveolin-1, focal adhesion kinase and other proteins involved in cell-cell and cell-extracellular matrix interactions (Diaz-Valdivia *et al*, 2020; Fu *et al*, 2020; Wadham *et al*, 2003; Zhang *et al*, 2012). The molecular mechanisms of its phosphatase activity are not clearly understood since its crystal structure and *in vitro* assays of PTPN14 biochemical activity suggest that it is an inactive phosphatase (Barr *et al*, 2006; Barr *et al*, 2009). Interestingly, in epithelial cells, independent of its phosphatase activity, PTPN14 regulates the Hippo–YAP mechanotransduction pathway. PPxY motifs in PTPN14 bind to WW motifs in pYAP1, leading to cytoplasmic sequestration of pYAP1 and consequent suppression of cell growth (Huang *et al*, 2013; Liu *et al*, 2013; Wang *et al*, 2012). Here we show that PTPN14 binds SMAD4, but in this case, it protects the latter from ubiquitination and protein turnover. In contrast, TAZ, the predominant YAP1 paralogue in primary human arterial ECs, is turned over by PTPN14. We therefore show a physical link between PTPN14 and SMAD4 that potentiates basal levels of SMAD-driven transcriptional activity downstream of ALK1 to support vascular quiescence while suppressing TAZ mediated signal transduction.

## RESULTS

### *Ptpn14* expression in adult lung associates with markers of angiogenesis and vascular remodeling, Rho kinase and TGFβ/BMP signaling, and endocytic recycling

We previously found genetic association between SNPs within *PTPN14* and the presence of pulmonary AVMs in a large panel of HHT patients (Benzinou *et al*., 2012). To shed light on the functional role of *Ptpn14* in the lung, we undertook gene expression network analysis on bulk RNA samples isolated from 69 individual, highly genetically heterogeneous wild type mice. We generated this genetically divergent panel of RNAs by interbreeding two different mouse species, namely *Mus spretus* and *Mus musculus,* in an F1 backcross to *M. musculus* (**Figure 1A**). Since the cross is inter-species, it shows limited fertility (see Methods), but provides greater inter individual genetic heterogeneity than that observed within the human population. Due to genetic variation across the panel each gene shows a wide dynamic range of expression, permitting powerful expression network analysis (Kim *et al*, 2013; Letteboer *et al*, 2015; Quigley *et al*, 2016; Quigley *et al*, 2009).

**Figure 1:**
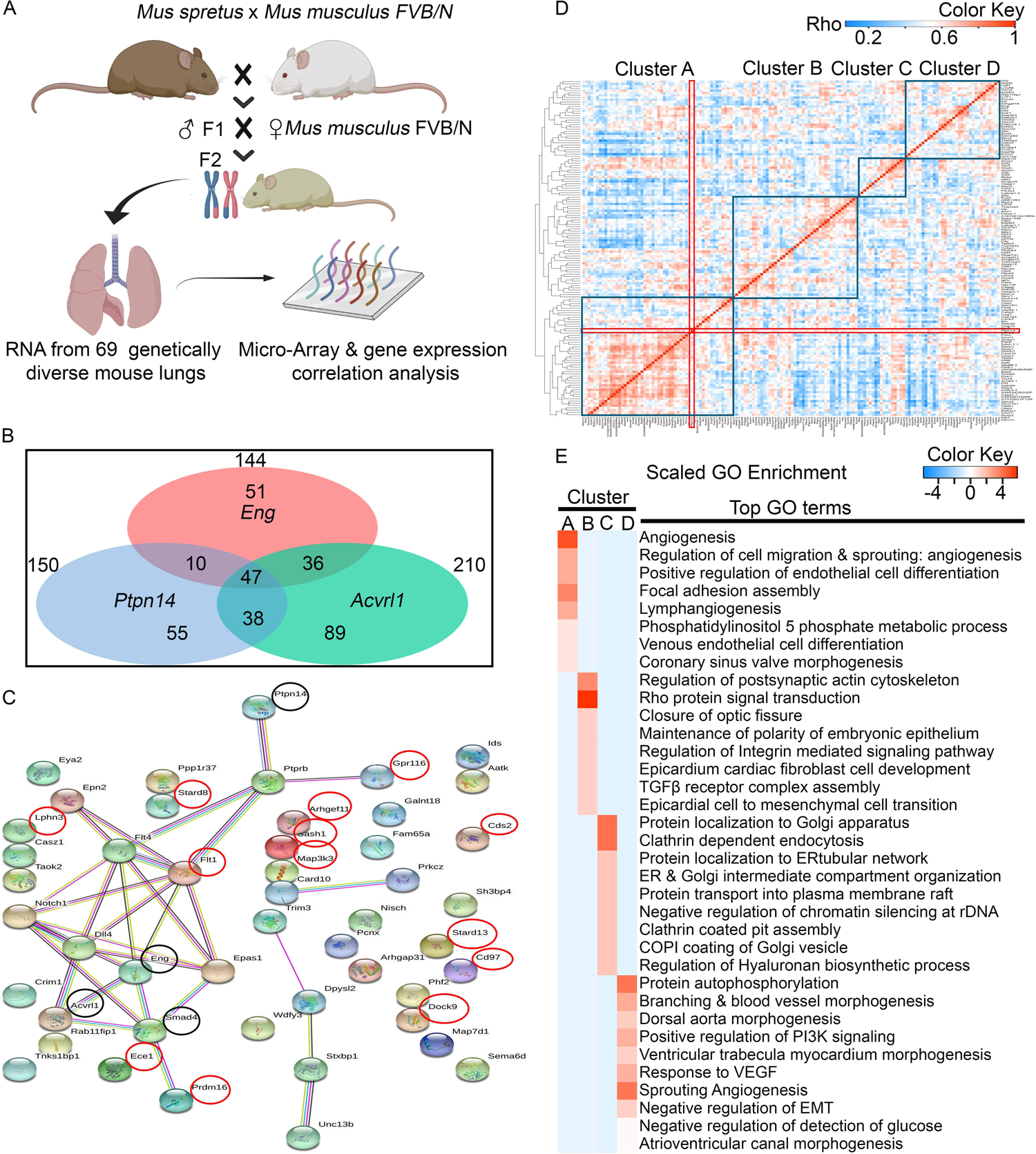
Unbiased gene expression network analysis of *Ptpn14* in adult lung implicates roles in angiogenesis and vascular remodeling, TGFβ/BMP and Rho kinase signaling, and endocytic recycling. A) Cartoon depicting the inter-specific backcross utilized (*Mus spretus* x *Mus musculus*). F2 mice were generated from backcrossing fertile female F1 hybrid mice to male *Mus musculus* FVB mice, and bulk RNAs harvested from individual lungs of 69 genetically distinct wildtype mice. **B)** Venn diagram shows overlap of *Ptpn14*-correlated transcripts with those correlated with *Acvrl1* and with *Eng* (at *rho* ≥ 0.54) across the 69-mouse lung panel. 47 genes show expression correlation with *Ptpn14, Eng,* and with *Acvrl1* in the lung (**Figure 1B**). (**C**) The 47 genes co-correlating with *Ptpn1, Eng and Acvrl1,* presented as an Ingenuity Pathways Network, that is based on preexisting knowledge of connectivity between genes. 13 of these 47 genes (circled in red), also correlate strongly with *Smad4* (*rho ≥ 0.54*). The four seed genes, *Ptpn14, Acvrl1, Eng,* and *Smad4*, are circled in black. D) Each of the 150 *Ptpn14-* associated genes underwent correlation analysis. Data underwent unsupervised hierarchical cluster analysis based on *rho* value for each pair-wise gene comparison, with the heat map generated indicating the magnitude of each pair-wise correlation coefficient. This reveals four gene clusters labelled A to D. Double red lines indicate position of *Ptpn14* gene in both axes. **E)** Gene Ontology (GO) analysis for each of the four clusters found in (**D).**

Across the panel of 69 mouse lung RNAs, 150 transcripts showed correlation with *Ptpn14* expression exceeding a threshold of *rho ≥ 0.54*; (*z score >3.5; P< 0.0003)*. These were enriched for genes involved in angiogenesis, cardiovascular morphogenesis, protein phosphorylation pathways, TGFβ/BMP signaling, rho kinase signaling, cell-cell interactions and endocytosis, as revealed by Gene Ontology (GO) analysis (**Supplementary Table S1**). This finding supports the notion that *PTPN14* is involved in regulation of BMP modulated vascular biology. Notably, 63% of the *Ptpn14-*correlated gene-set (95 of 150 genes) also exhibited strong expression correlation (*rho≥ 0.54*) with the endothelial-specific HHT causative genes, *Eng* or *Acvrl1* (**Figure 1B**), with a core of 47 *Ptpn14-*associated genes correlated with both *Eng* and *Acvrl* (**Figure 1B,C; Supplementary Table S2**), suggesting not only expression in the same cell type, but also in the same functional pathway (Quigley & Balmain, 2009; Quigley *et al*, 2011).

We then analyzed each of the 150 *Ptpn14-*associated genes for pair-wise expression correlation with each other, followed by unsupervised hierarchical cluster analysis based on the resultant *rho* value for each pair-wise comparison (**Figure 1D,E).** This revealed four gene clusters (**Figure 1D,E; Supplementary Table S3**). Cluster A (including *Ptpn14*, *Acvrl1*, *Notch1*, and *Amotl2*) showed enrichment for genes of angiogenesis, EC differentiation, focal adhesion, lymphangiogenesis and venous differentiation. Cluster D (including *Eng, Dll4, Map3k3, Flt1, and Flt4*) showed enrichment for genes involved in angiogenesis, cardiovascular morphogenesis, VEGF responses and PI3K signaling. Cluster B was enriched for Rho protein signaling, TGFβ receptor complex assembly, integrin mediated signaling and epicardial-to-mesenchymal cell transition, and Cluster C was enriched for processes of protein trafficking within the ER and Golgi apparatus, assembly of clathrin-coated pits and endocytosis (**Figure 1E**).

The third HHT-causative gene*, Smad4,* has a broader cell-specific expression distribution than *Eng* or *Acvrl1* thus its correlation coefficient with *Ptpn14,* although strong (*rho = 0.50*), was just below the threshold we had selected for the *Ptpn14-*correlated gene-set (*rho ≥ 0.54*) (**Figure 1B,D**). Nevertheless, of the 47 lung transcripts that correlated with *Eng, Acvrl1,* and *Ptpn14* (*rho > 0.54*), thirteen also showed correlations with *Smad4* (*at rho ≥ 0.54:* **Figure 1C).** Of these 13 genes, the strongest associations were with *Flt1,* encoding FLT/VEGFR1; *Ece1,* encoding endothelin-converting enzyme 1*; Sash1* and *Map3k3* (**Supplementary Table S4**). The latter two are reportedly involved in Hippo signaling (Jiang *et al*, 2020; Lu *et al*, 2021). *Map3k3* is additionally involved in stress-activated protein kinase signaling and regulation of NFkB activation (Ellinger-Ziegelbauer *et al*, 1997; Yang *et al*, 2001), and recent studies show interaction with TGFβ signaling in arterial remodelling (Deng *et al*, 2021a; Deng *et al*, 2021b). Eight of the other thirteen genes encode components of G protein coupled receptor (GPCR) signaling pathways related to Cdc42 rho kinase that influences cytoskeletal regulation, cell adhesion and migration (Sakabe *et al*, 2017; Yoshida *et al*, 2021). Overall, this powerful inter-specific gene expression network analysis provides strong support for PTPN14 interacting with HHT pathway components to regulate vascular cell adhesion, migration, and mechanotransduction, in lung in vivo.

### PTPN14 is expressed in lung and heart ECs in vivo

To validate that PTPN14 is expressed in ECs in vivo, we examined *Ptpn14* RNA and protein localization *in situ* in developing embryos and in adult lung and heart. Chromogenic RNAScope® *in situ* hybridization analysis revealed expression of *Ptpn14 RNA* (blue) within *Eng-*positive (pink) blood islands and vessels of the 8.5 days *post-coitum* yolk sac as well as in *Eng-*negative embryonic neuroepithelium (**Figure 2A, B**). In adult lung, *Ptpn14* RNA was expressed along with *Eng* in ECs of blood vessels and alveoli, as well as in *Eng-*negative bronchial epithelial cells (**Figure 2C, D**). Preferential expression of *Ptpn14* in ECs of adult lung is supported by analysis of single cell RNA-seq data (Endale *et al*, 2017; Travaglini *et al*, 2020) where ECs express highest *Ptpn14* RNA levels, with sequentially lower expression in mesenchymal, epithelial, and immune cells of both mouse and human lung (**Supplementary Figure S1**). In the heart and great vessels *Ptpn14* and *Eng* transcripts were co-expressed in endocardial cells and in ECs, as illustrated in the left cardiac atrium and aorta, whereas in the ventricular myocardium and interventricular septum of the heart, *Ptpn14* was expressed in both in cardiomyocytes and ECs of the coronary capillaries (**Figure 2E-G**). *RNAScope* analysis using non-specific probes showed no signal (**Supplementary Figure S2**).

**Figure 2:**
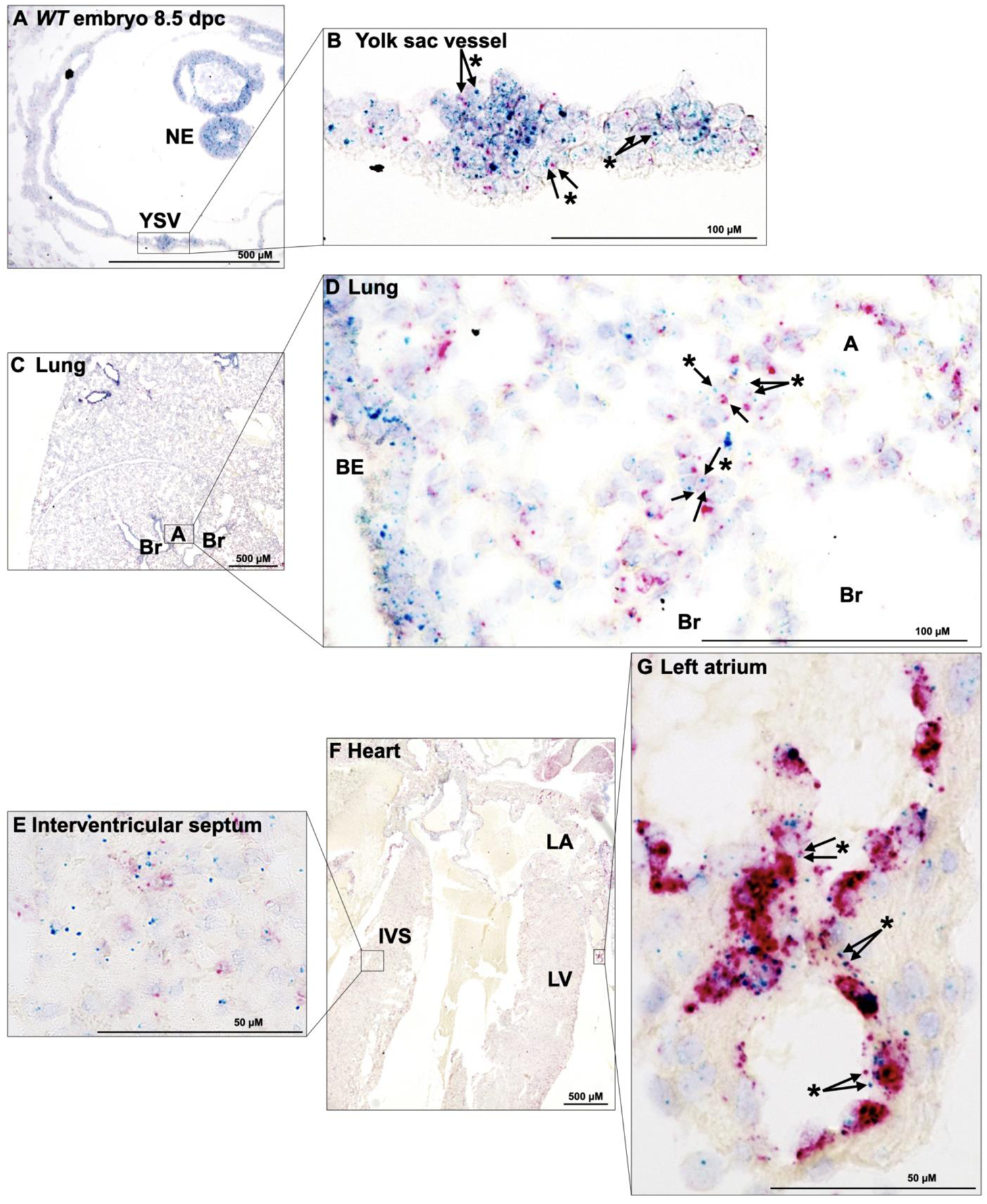
*Ptpn14* expression in ECs of multiple tissues. *Ptpn14* (cyan) and *Eng* (pink) RNA localization by *in situ* hybridization (RNAScope®) in tissues harvested from C57BL/6J mice. **A,B)** 8.5 dpc embryos; *Ptpn14* RNA localizes to *Eng*-negative embryonic neuroepithelium (NE) and *Eng*-positive extraembryonic yolk sac vessels and blood islands (YSV). **C,D**) In lung, *Ptpn14* and *Eng* co-localize in cells of the pulmonary blood vessels and alveoli (*), and *Ptpn14* but not *Eng* is expressed in bronchiolar epithelium (BE). (A), alveoli; (Br), bronchiole, (BE) bronchial epithelium). **E-G**) In heart, *Ptpn14* and *Eng* co-localize to endocardial cells (*) lining the left atrium (LA) and capillary ECs of the myocardium, such as within the interventricular septum (IVS) and left ventricle (LV). All tissues were counterstained with hematoxylin. Asterisks denote cells co-labeled with both *Eng* and *Ptpn14* probes.

Immunohistochemical analysis of PTPN14 protein was concordant with the distribution of *Ptpn14* RNA, showing extensive co-staining for PTPN14 and PECAM-1 proteins in ECs of pulmonary veins and alveolar capillaries (**Figure 3**; yellow-orange staining in composite view). Variable levels of PTPN14 protein staining were observed in PECAM-1-negative bronchial epithelia, with intense PTPN14 staining in Clara cell-rich epithelia of the terminal bronchioles, and lower expression primarily localized to basolateral regions of the simple cuboidal epithelium of the respiratory bronchioles (**Figure 3**). Smooth muscle cells underlying some bronchioles and blood vessels also showed PTPN14 staining (Green stain in composite exposure; **Figure 3**).

**Figure 3.**
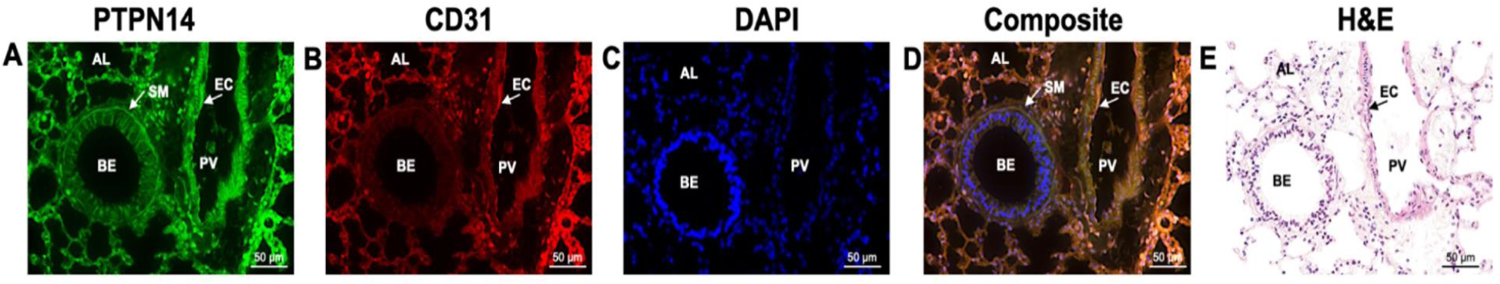
Detection of PTPN14 protein in PECAM-1 positive ECs of lung in vivo. **A-D representative** 5uM section of mouse lung, showing staining for (**A**) PTPN14 (green); (**B**) PECAM-1/CD31 (red); and (**C**) DAPI (blue). D, shows yellow to orange staining in composite exposure indicating co-expression of PTPN14 with the endothelial marker, CD31. (**E**) H and E staining of an adjacent section, showing. AL, alveoli; BE, bronchial epithelium; EC, endothelial cell lining pulmonary vein; PV, pulmonary vein; SMC, smooth muscle cell.

### PTPN14 supports endoglin, ALK1 and SMAD4 protein expression in primary human ECs

To investigate possible interactions between PTPN14 and the BMP9/ALK1 signaling pathway in primary human ECs, we examined the effects of *siRNA*-mediated knockdown (KD) of *ENG (siENG)*, *ACVRL1 (siALK1)*, and *PTPN14* (*siP*) on BMP9 signaling in low passage primary human umbilical arterial ECs (HUAECs). Because *PTPN14* expression is highest at high cell density (Wang *et al*., 2012) (**Figure S3A**), HUAECs were transfected with *siRNAs* and cultured to >90% density before treatment with BMP9 to mimic conditions of a stable vascular endothelial monolayer.

In control HUAECs transfected with non-targeting *siRNA* (*siNT*), BMP9 treatment led to transient phosphorylation of SMAD1/5, and marginally elevated PTPN14 levels. This activation of R-SMADs 1 and 5 was eliminated by prior KD of ALK1 (*siALK1)*, a SMAD1/5 phosphorylating receptor kinase (**Figure 4A**). 24 hour exposure of primary HUAECs to BMP9 induced endoglin levels, and this was lowered or ablated by transfection of *siALK1* (**Figure 4A**) or *siSMAD4* (**Figure S3B,C**) consistent with the encoding *ENG* gene being a known BMP transcriptional target (Calvo-Sánchez *et al*., 2019; Morikawa *et al*., 2011; Tual-Chalot *et al*., 2014). Contrasting with *siALK1*, *siENG* did not fully suppress BMP9-induced SMAD1/5 phosphorylation (**Figure 4A**), suggesting that endoglin is not essential for BMP9-induced activation of canonical SMAD1/5 signaling, or it is required at only low levels, consistent with observations of others (Baeyens *et al*, 2016; Upton *et al*, 2009) and suggesting that in arterial ECs, endoglin is a predominantly a target of BMP9/ALK1 signaling rather than a major upstream mediator of signaling.

**Figure 4:**
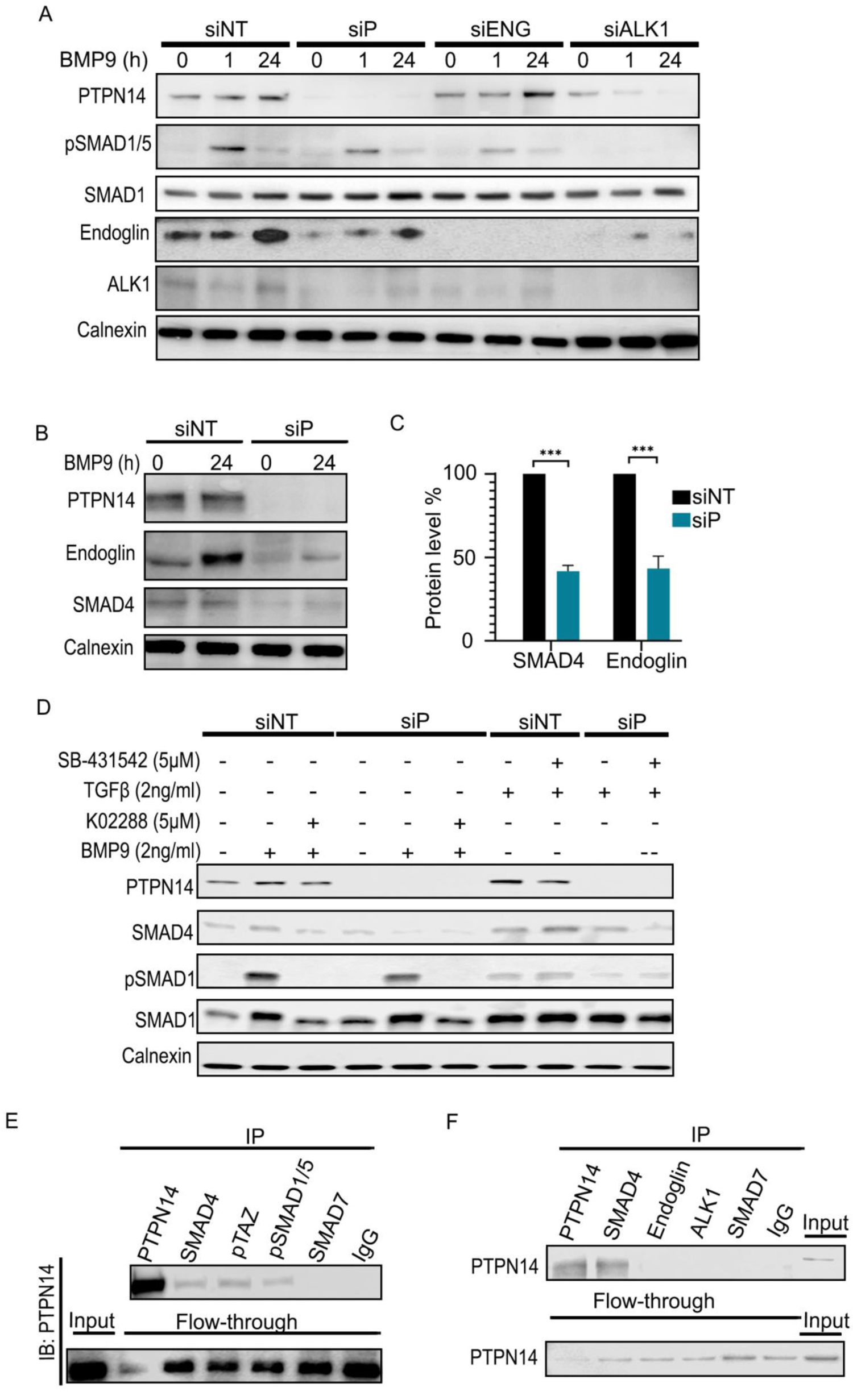
Reciprocal effects of *PTPN14* expression on HHT signaling components in primary human arterial ECs. HUAECs were transfected with the specified *siRNAs.* 48 hours later, protein lysates were harvested following treatment with BMP9 (2ng/ml) for 0, 1 or 24 hours, as indicated. (**A-B**) Representative western blots (of n=3 biological replicates). **C)** Quantification of westerns showing SMAD4 protein levels **in siSMAD4 transduced cells** relative to non-targeting, *siNT,* transfected HUAECs averaged from three biological replicates. Examples of replicates and the use of different *siRNAs* are shown in **Supplementary Figure S3**. **D)** Western blot analysis of protein lysates from *siRNA*-treated HUAEC lysates after 1h treatment with TGFβ (2ng/ml) or BMP9 (2ng/ml), with or without 5uM K02288 (an ALK1 inhibitor) or 5uM SB431542 (a TGFβR1 inhibitor). **E)** Co-immunoprecipitation using antibodies against endogenous PTPN14, SMAD4, pTAZ, pSMAD1/5, SMAD7, and control IgG, and probed for interaction with endogenous PTPN14. **F)** Co-immunoprecipitation using antibodies against endogenous PTPN14, SMAD4, endoglin, ALK1, SMAD7, or IgG controls, and probed for interaction with endogenous PTPN14.

*siPTPN14* had a similar effect to *siENG* in reducing the induction of SMAD1/5 phosphorylation by BMP9 (**Figure 4A**), and we found that *siPTPN14* also lowered steady state SMAD4 levels (**Figure 4B-C**). Consistent with this reduction in pSMAD1/5 and SMAD4 levels, *siPTPN14* reduced basal and BMP9-induced endoglin levels (**Figure 4A-C**). Intriguingly, in contrast to the effects of *siALK1* in suppressing BMP9-induced PTPN14 expression*, siSMAD4* and *siENG* both elevated PTPN14 levels, especially after BMP9 treatment (**Supplementary Figure S3C**), suggesting a complex interplay of negative and positive feedbacks.

Treatment with TGFβ1 induced phosphorylation of SMAD1 in HUAECs, but the magnitude of this activation was far less than that observed in response to BMP9 (**Figure 4D**). Since SMAD1 activation by TGFβ was resistant to treatment with the TGFβRI (ALK5) inhibitor, SB431542, we conclude that its effects are likely mediated through TGFβR2 activation of ALK1 kinase (Goumans *et al*, 2003) (**Figure 4E**). Indeed, SB431542 potentiated total SMAD1 and SMAD4 levels in the presence of TGFβ, indicating that when TGFβR1 kinase is inhibited, TGFβR2 signaling is diverted down the ALK1-SMAD1/5/SMAD4 pathway (Goumans *et al*, 2002). KD of *PTPN14* reduced SMAD4 levels and the induction of SMAD1 phosphorylation by both BMP9 and TGFβ1 (**Figure 4D**), suggesting a role for PTPN14 in stabilizing pSMAD1/SMAD4, regardless of the activating ligand or type II receptor.

### Endogenous PTPN14 binds SMAD4 but not endoglin or ALK1 in primary HUAECs

Since PTPN14 regulates endoglin, SMAD4 and ALK1 protein levels, we tested the possibility that PTPN14 might directly bind to at least one of these proteins. As a positive control, we tested YAP1, a well characterized binding partner of PTPN14 in epithelial cells (Huang *et al*., 2013; Liu *et al*., 2013; Wang *et al*., 2012), but this protein was undetectable in high density primary HUAECs. We therefore tested for interaction between PTPN14 and the YAP1 paralogue, TAZ, which is robustly expressed in quiescent HUAECs. As previously reported for pYAP1/PTPN14 interaction in epithelial cells, PTPN14 co-immunoprecipitated with phosphorylated (p)TAZ in ECs (**Figure 4E**), which is the first documented observation of physical interaction between pTAZ and PTPN14. Endogenous SMAD4 and pSMAD1/5 also co-immunoprecipitated with PTPN14, and the magnitude of this binding was like that between PTPN14 and pTAZ (**Figure 4F**). Notably SMAD7, an inhibitory SMAD, did not co-immunoprecipitate with PTPN14 (**Figure 4E,F),** nor did ALK1 or endoglin (**Figure 4F**). Interaction between PTPN14 and the BMP9 signaling pathway therefore appears to be at the level of SMAD proteins, downstream of endoglin and the ALK1 receptor.

### Endogenous SMAD4-PTPN14 interactions in HUAECs detected by proximity ligation assay in situ

To confirm the presence of molecular interactions between endogenous PTPN14 and SMAD4 proteins in primary HUAECs, we undertook proximity ligation assays (PLAs) in low passage HUAECs at high cell density, using primary antibodies optimized for specificity to each target protein as assessed by immunofluorescence following *siRNA* KD (**Supplementary Figure S2B)**. Using a PCR-based signal initiated by primers that tag distinct secondary antibodies, PLA detects proteins within 40nM of each other within the cell, i.e. proteins presumed to bind one another. As a positive control, we undertook PLA between SMAD4 and pSMAD1/5. These showed PLA signals in both nucleus and cytoplasm and this interaction signal increased after BMP9 treatment for 1 hour (**Figure 5A-F**). PLA analysis for interaction between PTPN14 and SMAD4 also showed nuclear and cytoplasmic signals (**Figure 5P, Supplementary Figure S5**), and this interaction was enhanced in both sub-cellular compartments one hour after BMP9 treatment (**Figure 5G-L**). Control PLA reactions, undertaken using only one primary antibody or both secondary antibodies without the corresponding primaries, showed little to no background signal (**Figure 5M-O**). In conclusion, we demonstrate that physical interactions between SMAD4 and PTPN14 and between SMAD4 and pSMAD1/5 occur within both cytoplasm and nucleus of primary HUAECs, and the number of molecular interactions increases in both nucleus and cytoplasm after 1 hour of BMP9 treatment.

**Figure 5:**
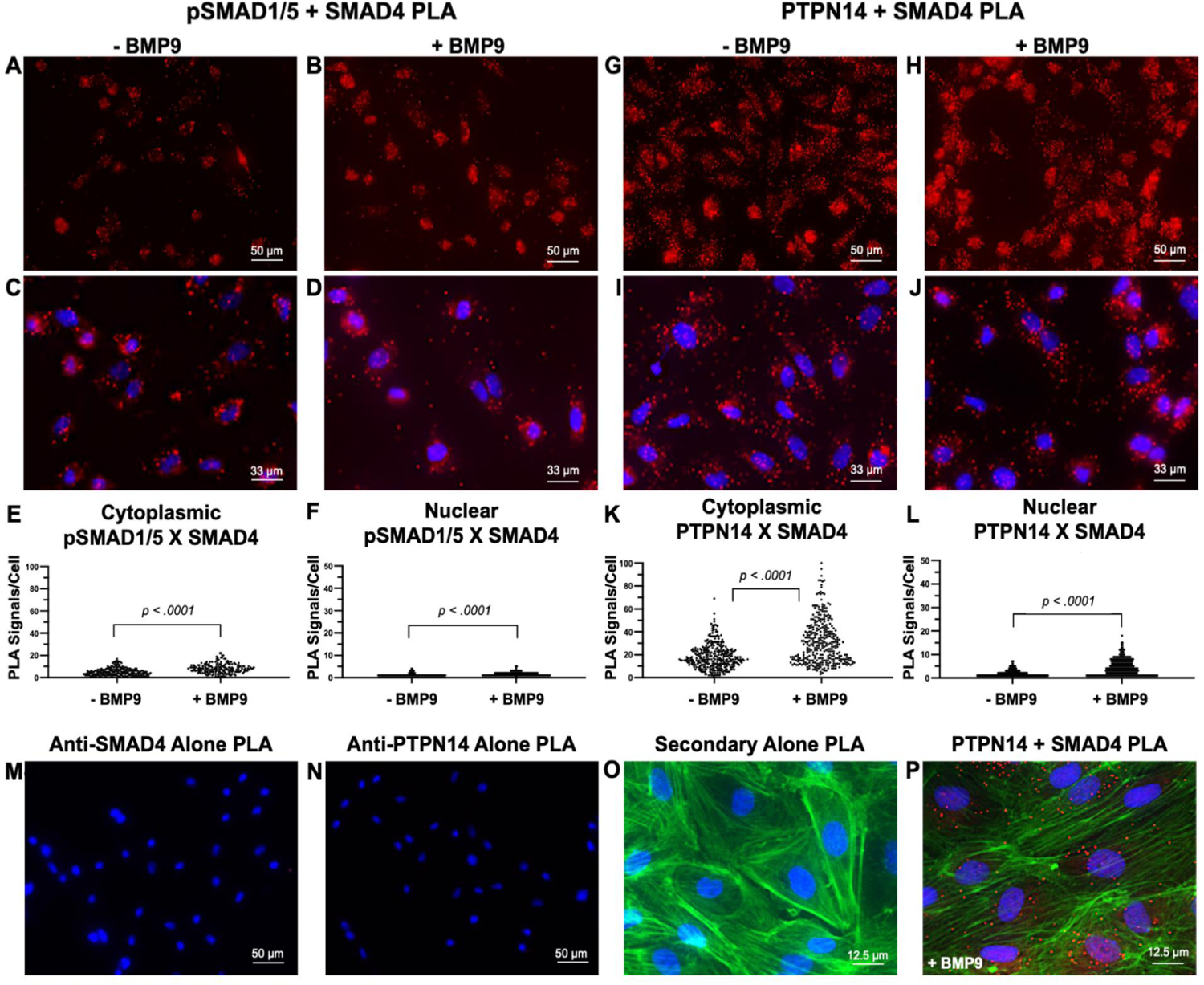
Molecular interactions between endogenous PTPN14 and SMAD4 proteins detected by PLA in HUAECs. PLA was undertaken to investigate physical interactions between endogenous SMAD4 and PTPN14 proteins within low passage HUAECs. Cells were pretreated or not for 60 minutes with 2ng/ml BMP9 prior to the PLA (as indicated). PLA signal (red) indicates molecular interaction between anti-pSMAD1/5 and anti-SMAD4 antibodies (**A-D**) as quantified according to nuclear and cytoplasmic signals in **E** and **F,** or between anti-PTPN14 with anti-SMAD4 antibodies (**G-J**), as quantified according to nuclear and cytoplasmic signals in **K** and **L**. Note that the nuclear to cytoplasmic ratio of the interaction signals did not vary after BMP9 treatment. Nuclei were counterstained using DAPI (blue). **M-O**) show examples of negative controls for PLA, undertaken with one primary antibody only or with two secondary antibodies only, as indicated. In **O** and **P**, cells were stained with Phalloidin prior to the PLA assay to reveal stress fibers and cellular outline. Scale bars are indicated in each frame.

### PTPN14 stabilizes SMAD4 levels in the cytoplasm and nucleus without affecting BMP9-induced nuclear shuttling

Nucleocytoplasmic fractionation revealed that PTPN14 was equally distributed between nucleus and cytoplasm in high density primary HUAECs (**Fig 6A, B**), with total PTPN14 levels reduced at low cell density (**Supplementary Figure S3A**) (Wang *et al*., 2012). SMAD4 was also found in both the nuclear and cytoplasmic fractions, with higher nuclear levels under the high cell density culture conditions used. Nuclear SMAD4 protein was consistently lowered by *siPTPN14*, whereas the effect of *siPTPN14* on cytoplasmic SMAD4 was difficult to assess, in part due to lower levels of cytoplasmic SMAD4 (**Figure 6A, B**). Despite reduced SMAD4 expression following *siPTPN14* treatment, nuclear accumulation of SMAD4 in response to BMP9 treatment still occurred (**Figure 6B)**, suggesting that PTPN14 is not required for nuclear shuttling of SMADs but is rather involved in stabilizing total SMAD4.

**Figure 6:**
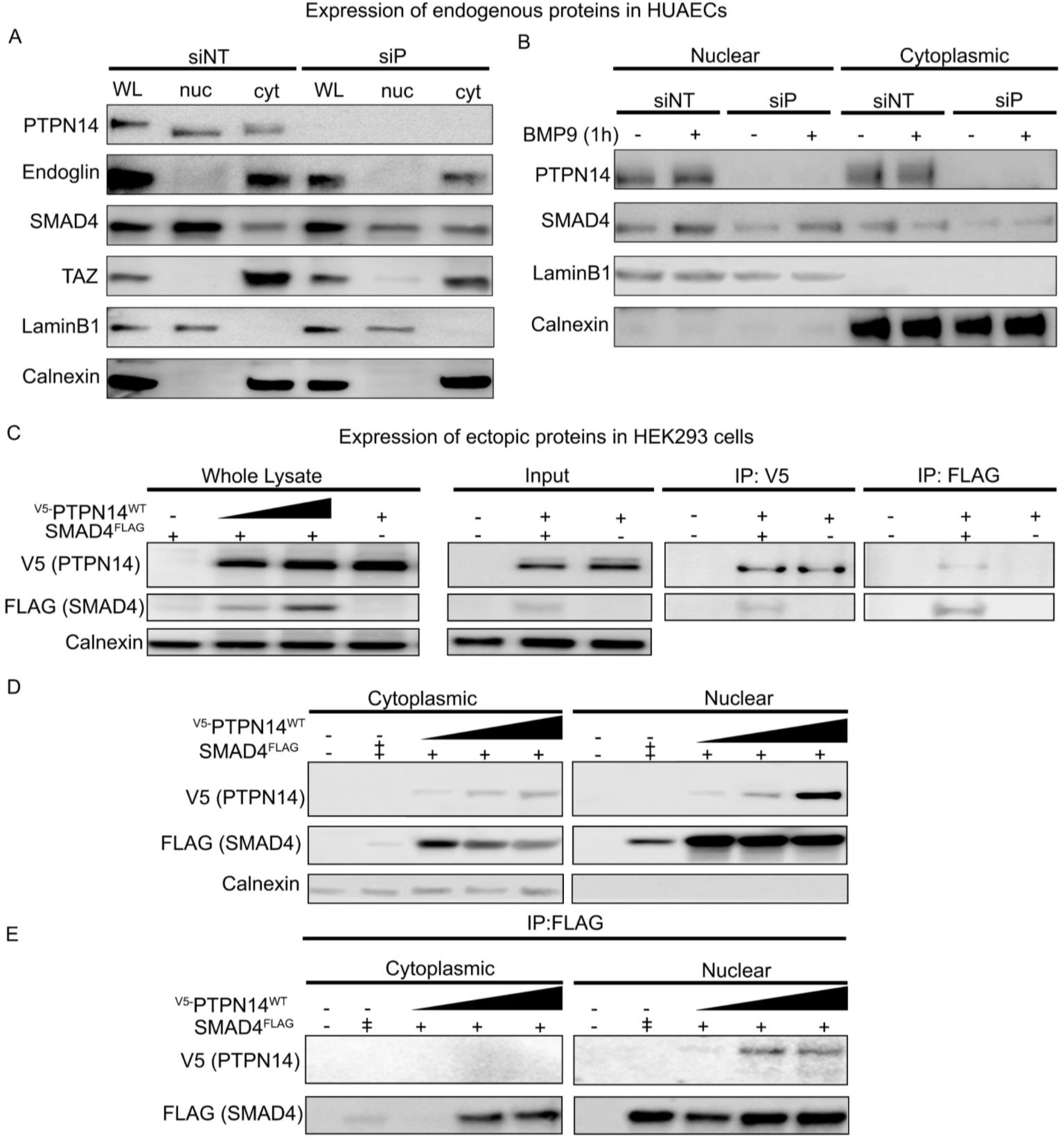
PTPN14 binds and regulates SMAD4 levels. **A)** Western blot analysis of protein lysates from *siRNA*-transfected HUAECs (no BMP9), showing effects of *siPTPN14* on endoglin, SMAD4 and TAZ protein levels in whole cell lysates (WL) or in fractionated nuclear (nuc) and cytoplasmic (cyt) lysates. **B)** Western blot of *siRNA*-transfected HUAECs with or without BMP9 treatment for 1h, showing effects on SMAD4 levels in nuclear compared to cytoplasmic lysates. **C)** Western blot and reciprocal co-immunoprecipitation analysis of ectopically expressed proteins after transient transfection into HEK293 cells. FLAG-SMAD4 was co-transfected with increasing quantities of V5-PTPN14, followed by visualization with anti-V5 and anti-FLAG antibodies. Note, stabilization of FLAG-SMAD4 protein by PTPN14 (left panel) and physical association between the two proteins (right panels). **D)** Western blot analysis of cytoplasmic and nuclear lysates of HEK293 cells after co-transfection of increasing concentrations of V5-PTPN14 and indicated quantities of FLAG-SMAD4, showing stabilization of SMAD4 predominantly in the nuclear fraction. **E)** Co-immuno-precipitation of HEK293 cytoplasmic and nuclear lysates (left to right) with an anti-FLAG antibody after co-transfection of V5-PTPN14 and FLAG-SMAD4, showing physical interaction between two proteins occurs predominantly within the nucleus. Calnexin and LaminB1 are cytoplasmic and nuclear markers, respectively.

To further investigate the relationship between PTPN14 and SMAD4, we overexpressed these proteins in HEK293 cells. Notably, co-transfection of FLAG-SMAD4 together with increasing quantities of V5-PTPN14 indicated that V5-PTPN14 stabilizes the FLAG-SMAD4 protein in a dose-dependent manner (**Figure 6C,** left panel). Physical interaction between V5-PTPN14 and FLAG-SMAD4 was detected by reciprocal co-immuno-precipitation (**Figure 6C,** three right panels), supporting our observation of interactions between the endogenous proteins in HUAECs (**Figures 4E, F, and 5**).

Subcellular fractionation of transfected HEK293 cells showed that FLAG-SMAD4 localizes to both nucleus and cytoplasm, and co-transfection with V5-PTPN14 raised FLAG-SMAD4 levels in both cellular compartments (**Figure 6D**). V5-PTPN14 and FLAG-SMAD4 interaction was observed primarily in the nucleus by co-immunoprecipitation (**Figure 6E**). The inability to detect such an interaction in the cytoplasm compared to PLA findings of endogenous interactions in HUAECs, may be due to low cytoplasmic FLAG-SMAD4 expression levels in HEK293 cells (**Figure 6D**), and/or use of different cell types and detection systems.

### FLAG-SMAD4 protein is stabilized by PTPN14 in HEK293 cells independent of its action on YAP

PTPN14 protein has three structural domains, a ∼ 300 aa amino terminal FERM domain that is characteristic of proteins associated with the cytoskeletal cortex (aa 21 to 306), a carboxy-terminal tyrosine phosphatase domain (aa 909 to 1180), and a long proline-rich linker region that spans between these and encompasses at least two PPxY motifs (**Figure 7A**). PPPY at aa 566-569 and PPEY at aa 748-752 are reported to bind phosphorylated YAP1 in epithelial cells, tethering it within the cytoplasm for ubiquitination and subsequent degradation via the proteasome (Huang *et al*., 2013; Liu *et al*., 2013; Wang *et al*., 2012). Our previous studies in HEK293 cells show that co-transfection of wild type V5-PTPN14 does not affect total YAP1 levels but shifts YAP1 localization from nucleus to cytoplasm to reduce YAP1/TEAD-driven transcription (Liu *et al*., 2013) consequent to cytoplasmic sequestration of pYAP1 by PTPN14. This cytoplasmic sequestration was independent of the FERM and phosphatase domains of PTPN14, but dependent on pYAP1 binding to the two PPXY motifs within the PTPN14 linker region (Liu *et al*., 2013).

**Figure 7:**
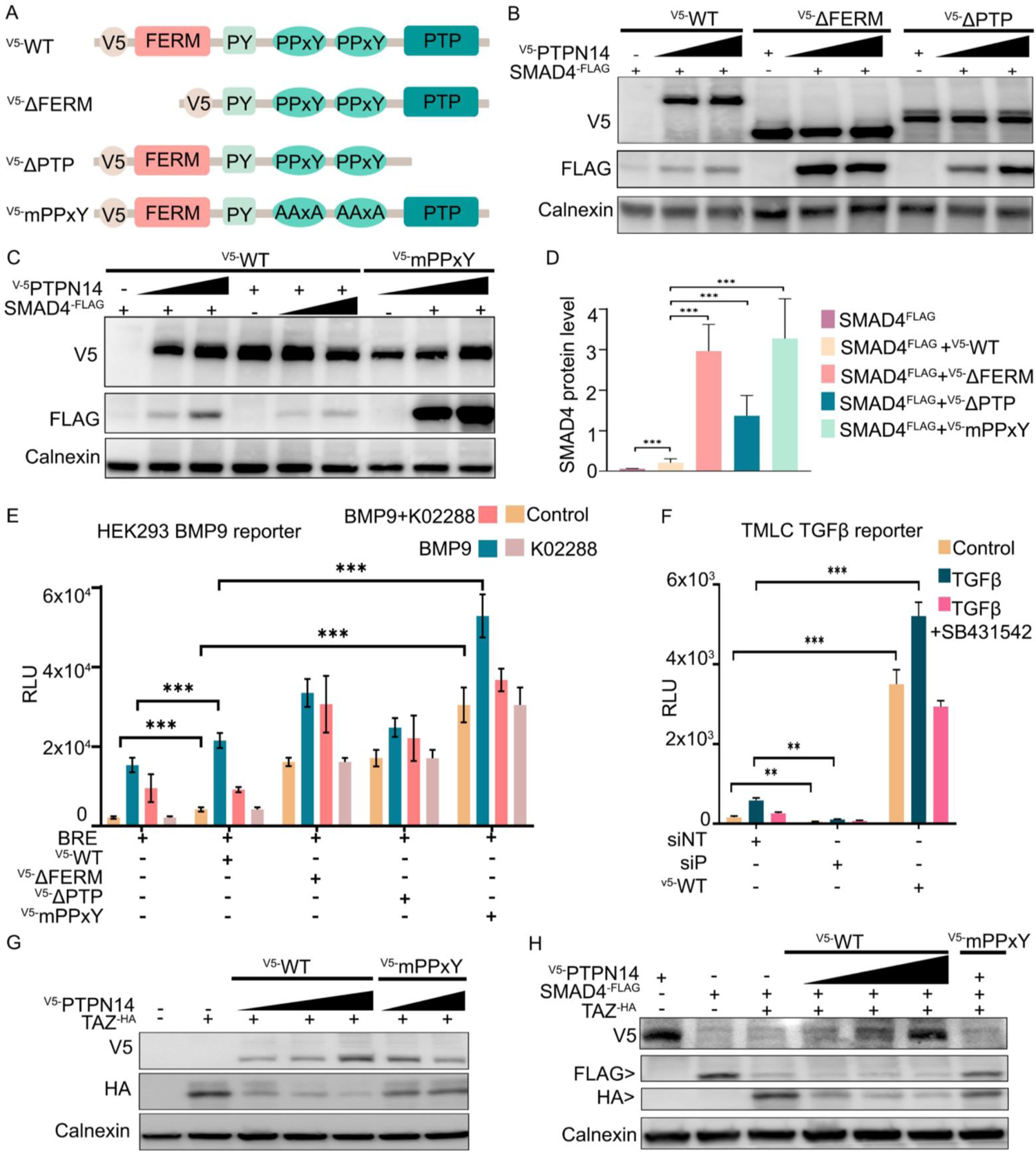
PTPN14 stabilizes SMAD4 protein and enhances SMAD4-dependent transcriptional readouts. **A)** Schematic representation of PTPN14 protein primary structure indicating different V5-PTPN14 mutant expression constructs used. **B-C)** Western Blot analysis of HEK293 cell lysates after co-transfection with FLAG-SMAD4 and increasing levels of the different V5-PTPN14 constructs (PTPN14^WT^, ΔFERM, mPPxY, and ΔPTP). **D)** Quantification of relative FLAG-SMAD4 protein levels after co-transfection with indicated PTPN14 mutants (data are the average of 3 biological replicates). **E)** Representative plots of BRE-Luciferase reporter assays in HEK293 to show the effect of different V5-PTPN14 constructs on basal and BMP9-induced luciferase-expression in the presence or absence of the ALK1 inhibitor, K02288. **F)** TGFβ-responsive luciferase reporter assay in TMLCs showing the effect of manipulating PTPN14 expression on basal and TGFβ-induced luciferase-expression, in the presence or absence of the TGFβR1 receptor inhibitor, SB431542. **G)** Western blot analysis showing the effects of V5-PTPN14 wild type and mPPxY.PTPN14 expression on co-transfected TAZ-HA levels. **H)** Western blot analysis of HEK293 cells co-transfected with V5-PTPN14 wild type or mPPxY.PTPN14, with or without FLAG-SMAD4 and/or TAZ-HA. ** *p* ≤ 0.01; *** *p* ≤ 0.001.

To determine the domain-specific contributions of PTPN14 to stability of SMAD4, we transfected V5-PTPN14 mutant expression constructs (**Figure 7A** (Liu *et al*., 2013)), together with FLAG-SMAD4, into HEK293 cells. In contrast to the negative effect of PTPN14 on YAP1 activity (Liu *et al*., 2013), we found that wild type V5-PTPN14 elevated FLAG-SMAD4 levels in a dose dependent manner (**Figure 6C and 7B-D**). V5-PTPN14-△FERM which lacks the PTPN14 ubiquitination site(s) (Wang *et al*., 2012), and thus reduces PTPN14 turnover (Liu *et al*., 2013), dramatically elevated FLAG-SMAD4 levels over that potentiated by wild type V5-PTPN14, as did the V5-PTPN14-ΔPase mutant (**Figure 7B**), indicating that neither the FERM nor phosphatase domains of PTPN14 are required for SMAD4 stabilization.

Moreover, this suggests that the PTPN14 proline-rich linker region is critical for SMAD stabilization. Transfection of the mPPxY.PTPN14 that harbors point mutations in the two YAP1-binding PPxY motifs, lead to the highest levels of co-transfected FLAG-SMAD4 protein, despite equivalent levels of V5-PTPN14 protein expression from the PTPN14 wildtype and mPPxY mutant forms (**Figure 7C**). Thus, unlike its effect on YAP1, PTPN14 potentiated higher FLAG-SMAD4 protein levels, regardless of the ability of PTPN14 to bind and sequester inactivate pYAP1 in the cytoplasm.

### PTPN14 potentiates transcriptional activity of BMP- and TGFβ-responsive reporters

To determine whether PTPN14-mediated stabilization of SMAD4 protein translates into potentiation of pSMAD1/5/4 mediated transcriptional responses, we examined the effects of PTPN14 overexpression on basal and BMP9 inducible transcription of a BMP-response element (BRE)-luciferase reporter in HEK293 cells. ALK1 is expressed endogenously in HEK293 cells (**Supplementary Figure S4A**), and BMP9 treatment for 24 hours induced BRE-luciferase reporter activity six-fold (**Figure 7E**). Co-transfection of BRE-luciferase with wildtype V5-PTPN14 significantly enhanced both basal and BMP9-induced BRE-luciferase activity. Each PTPN14 mutant construct that potentiated SMAD4 protein levels (**Figure 7B-D**) elevated basal BRE-reporter activity without potentiating the magnitude of luciferase induction by BMP9 (**Figure 7E**). Notably, the increase in basal BRE-luciferase activity by each PTPN14 construct was resistant to the ALK1 inhibitor, K02288 (**Figure 7E**), implying that it is independent of ALK1 kinase activity, consistent with the concept that enhancement of SMAD4 protein levels by PTPN14 occurs downstream of ALK1, bypassing a requirement for BMP9-induced ALK1 kinase activity to drive transcription of the BRE-luciferase reporter, and consistent with SMAD4 being a limiting factor in this transcriptional activity.

Since SMAD4 is a common co-SMAD used by all members of the TGFβ superfamily, we asked whether signaling via TGFβ-SMAD2/3 was also affected by PTPN14 expression. Indeed, *PTPN14* KD significantly reduced tonic and TGFβ-inducible expression of a PAI-1-luciferase reporter in TMLC cells (**Figure 7F**). Conversely, transfection of TMLC cells with WT-PTPN14 significantly enhanced basal PAI-I-reporter activity, and this was resistant to the TGFβR1 inhibitor, SB431542 (**Figure 7F)**. Elevated SMAD4 levels observed following PTPN14 over-expression therefore translate into enhanced tonic activity of both the BMP-responsive and TGFβ-responsive reporters with no increase in the magnitude of ligand inducibility. Nevertheless, relatively small modulations in SMAD4 levels by PTPN14 are transcriptionally and/or translationally amplified such that they significantly alter the signaling outputs. Following on from this observation, such changes might suppress or accentuate clinical phenotypes in a pathological setting. Notably, the *Tgfbm2* locus that harbors *Ptpn14* was originally identified as a suppressor of *Tgfb1-/-* mouse embryo lethality, and our current data are consistent with *Ptpn14* modulating both BMP9 and TGFβ signaling (Benzinou *et al*., 2012).

### PTPN14 interactions with SMAD4 and TAZ in HEK293 cells

In HUAECs, we found far higher expression of TAZ than YAP1. These two proteins are paralogues, however they have distinct protein structures and relatively little is known about PTPN14/TAZ interactions. To fill this knowledge gap, we undertook co-transfection experiments using V5-PTPN14 and HA-tagged TAZ (TAZ-HA) into HEK293 cells and asked whether these interactions affect the levels of co-transfected FLAG-SMAD4.

Wild type V5-PTPN14 decreased total TAZ-HA protein levels (**Figure 7G lanes 2-5)** and *siPTPN14* slightly potentiated expression of TAZ-HA in HEK293 cells (**Supplementary Figure S4B,C)**. This effect of wildtype V5-PTPN14 on total TAZ protein turnover contrasts with its effect on YAP1, where total levels of YAP1 are unaffected, but the nuclear to cytoplasm ratio is diminished by wild type V5-PTPN14 (Huang *et al*., 2013; Liu *et al*., 2013; Wang *et al*., 2012). Overexpression of V5-PTPN14.mPPxY was far less effective in reducing TAZ-HA levels than wild type PTPN14 (**Figure 7G, lanes 6-7**), concordant with the reduced ability of this mutant to bind TAZ.

When TAZ-HA was co-transfected with FLAG-SMAD4, SMAD4 was degraded. (**Figure 7H,** compare lanes 2 and 3). Addition of wildtype V5-PTPN14 to this co-transfection destabilized TAZ-HA but did not restore FLAG-SMAD4 levels (**Figure 7H,** lanes 4-6). Notably, the V5-PTPN14-mPPxY mutant that binds only weakly to TAZ failed to reduce either TAZ or SMAD4 levels (**Figure 7H, lane 7**). This might result if PTPN14 has overlapping binding domains for the two proteins such that TAZ competes with SMAD4. If SMAD4 binds within the proline-rich linker region of PTPN14 independent of the two PPxY motifs, then TAZ, bound to the PTPN14 PPxY motifs, might exert stearic hindrance on SMAD4/PTPN14 interaction. TAZ would outcompete SMAD4 for binding WT-PTPN14, resulting in SMAD4 degradation. The mPPxY.PTPN14 mutant however, which binds weakly to TAZ, is free to bind and stabilize SMAD4. In support of this concept, co-immunoprecipitation studies show that, while FLAG-SMAD4 binds all the mutant forms of PTPN14, it binds most avidly to the mPPXY.PTPN14 form (**Supplementary Figure S4D**).

### PTPN14 protects SMAD4 from ubiquitination and degradation

Our co-transfection experiments suggest that FLAG-SMAD4 protein is stabilized by PTPN14. We postulated that this may be due to protection of SMAD4 from ubiquitination and subsequent proteasomal degradation. To investigate this possibility, we probed the consequences of over-expression or *KD* of *PTPN14* on the ubiquitination state of SMAD4 and PTPN14. As expected, total SMAD4 levels were decreased by *siPTPN14* and increased by PTPN14 overexpression (**Figure 8A**). However, whereas PTPN14 overexpression resulted in higher total SMAD4 levels, the fraction of SMAD4 that was ubiquitinated was reduced (**Figure 8B**), despite increased ubiquitination of the over-expressed PTPN14 protein *per se* (**Figure 8C**). In contrast, though *siPTPN14* reduced total SMAD4 levels (**Figure 8A**), it did not result in an equivalent reduction in SMAD4 ubiquitination (**Figure 8B**), suggesting an absolute increase in the fraction of SMAD4 that was ubiquitinated.

**Figure 8:**
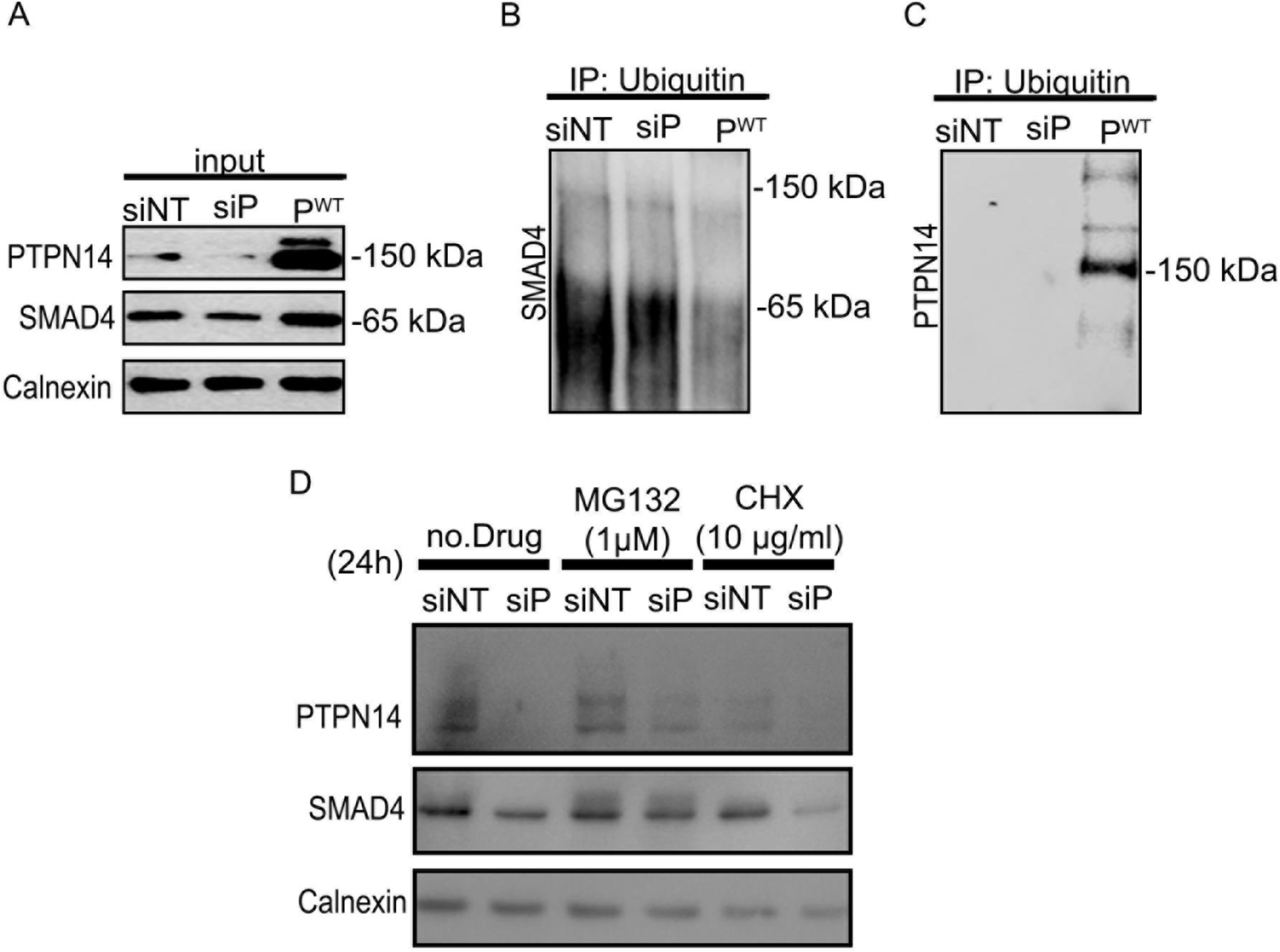
PTPN14 protects SMAD4 protein from ubiquitination and degradation. **A)** Western-blot analysis of HUAEC protein lysates after transfection with *siPTPN14* or V5-PTPN14^-WT^ compared to controls transfected with non-specific *siRNA, siNT. siPTPN14* decreases and PTPN14-WT increases total SMAD4 levels. (**B,C**) HUAEC protein lysates from (**A**) were immunoprecipitated using anti-ubiquitin antibodies, and probed for (**B**) SMAD4 and (**C**) PTPN14. **D)** Western blot analysis of SMAD4 after transfection of HUAECs with *siPTPN14* or *siNT* followed by 24h treatment with MG132 (1µM), cycloheximide (10 µg/ml), or no drug. Diminution of SMAD4 level by *siPTPN14* was rescued by proteasome inhibition (MG132), whereas cycloheximide potentiated this decrease, implying fast SMAD4 turnover in the absence of PTPN14.

We also examined the turnover of total SMAD4 protein in the presence and absence of PTPN14, by examining the effects of inhibitors of protein translation (cycloheximide) or proteasomal degradation (MG132). Whereas *siPTPN14* reduced total SMAD4 protein levels, this was prevented by MG132, but potentiated by cycloheximide treatment. Taken together, these data indicate that SMAD4 protein undergoes rapid turnover in human primary ECs, and PTPN14 stabilizes SMAD4 by prevention of its ubiquitination and subsequent turnover through proteasomal degradation.

## Discussion

We previously identified *PTPN14* as a gene harboring genetic variants that associate with pulmonary AVMs in type I and type 2 HHT patients (Benzinou *et al*., 2012). However, human genetic association data is only a first step in interrogating how a candidate gene might influence a human disease process. A molecular understanding of how the gene product regulates the pathobiology of the specific disease and how its expression interacts within gene networks to regulate in vivo biology may provide clues towards potential therapeutic approaches. Here we show, for the first time, by both co-immunoprecipitation and PLA, that endogenous PTPN14 directly binds to and stabilizes SMAD4 in quiescent primary human arterial ECs. PTPN14 thus physically interacts with this key component of the HHT signaling pathway in human primary ECs and protects endogenous SMAD4 from ubiquitination and subsequent degradation. We show that PTPN14 enhances tonic transcriptional output from both a BMP-responsive and a TGFβ-responsive reporter and, using specific inhibitors of these receptor kinases, we find that this is independent of signaling from the ALK1 or type I TGFβ receptor. Therefore, PTPN14 acts downstream of the TGFβ/BMP receptors, at the level of SMAD stabilization, to potentiate SMAD-mediated transcriptional readouts, such as expression of *ENG* that encodes endoglin. This conclusion is supported by our co-immunoprecipitation and PLA data that reveal that the PTPN14-SMAD4 interaction occurs in both nucleus and cytoplasm. SMAD4 interaction is enhanced by BMP treatment in both cytoplasm and nucleus, but PTPN14 does not modulate cytoplasmic to nuclear shuttling of SMAD4. As SMAD signaling is reduced in HHT ECs (Fernandez *et al*., 2005), PTPN14 may buffer this diminution of ALK1-SMAD1/5-SMAD4 signaling to minimize disease progression, such as formation of telangiectasias and AVMs.

We also show that in confluent primary human arterial ECs, TAZ is the major paralogue of YAP/TAZ expressed and that like YAP1, TAZ binds to PTPN14. In contrast to SMAD4, however, TAZ protein is turned over by PTPN14. Thus, PTPN14 likely contributes to vascular quiescence and protection from AVM formation through suppression of nuclear TAZ/TEAD signaling whilst promoting active SMAD signaling at a juncture downstream of the ligand activated ALK1 receptor. PTPN14 therefore provides a physical link between SMAD4 and YAP/TAZ, orchestrating vascular stability through multiple mechanisms including sequestration and turnover of pYAP/TAZ in the cytoplasm and support of SMAD4-mediated transcription in the nucleus. Recent studies reveal that PTPN14 can additionally contribute to maintenance of endothelial integrity by dephosphorylation of VE-Cadherin (Fu *et al*., 2020), thus acting on multiple molecular pathways to orchestrate stabilization of the quiescent vascular endothelium.

Our mouse lung expression network analysis demonstrates the power of an inter-specific mouse backcross to interrogate possible in vivo functions of candidate genes. (*Lung expression dataset is available* @ *data <u>to be deposited in NCBI GEO database on acceptance for publication</u>*). We demonstrate that *Ptpn14* shows gene expression network interdependence with the major HHT causative genes and with multiple EC-specific markers in quiescent mouse lung in vivo. This was supported by RNA *in situ* hybridization analysis and immuno-histochemical staining, providing empirical evidence that *PTPN14* is expressed in ECs of the lung. Intriguingly our expression network analysis shows that 8 of 13 genes showing strong expression correlates with *Ptpn14, Eng, Acvrl1* and *Smad4*, encode components of GPCR signaling pathways, particularly implicating Cdc42 signaling. A small GTPase of the Rho family, Cdc42 regulates signaling pathways that control endocytosis, cell shape, morphology, and migration, and can be activated by integrin signaling. It is central to dynamic actin cytoskeletal assembly and rearrangement that forms the basis of cell-cell adhesion and migration. This has relevance to HHT molecular pathogenesis, since genetic loss of endoglin, ALK1, or SMAD4 results in changes in cell shape, cell size, and loss of the characteristic directional migration of ECs against blood flow (Crist *et al*, 2018; Jin *et al*., 2017; Ola *et al*, 2018; Poduri *et al*, 2017; Sugden *et al*., 2017). Given that PTPN14 modulates nuclear and cytoplasmic levels of YAP/TAZ, as well as TAZ turnover, it is intriguing that Cdc42 has been implicated in mediating the promigratory effect of cytoplasmic YAP/TAZ on endothelial tip cell migration in developing retina (Sakabe *et al*., 2017).

We find that expression of *Map3k3*, encoding Mitogen-Activated Protein Kinase Kinase Kinase 3 (MEKK3/MAP3K3) is tightly correlated with expression of our four seed genes (*Eng, Acvrl1, Smad4* and *Ptpn14*). MAP3K3 has been suggested to activate Hippo signaling at two junctures; by direct phosphorylation and activation of LATS1/2 and YAP1, and by promoting β-TRCP interaction with YAP1 to drive YAP1 ubiquitination and degradation (Lu *et al*., 2021). Significantly, MAP3K3 is a mediator of vascular lesion development in a related but distinct autosomal dominant disorder, Cerebral Cavernous Malformations, wherein haploinsufficiency for one of three CCM genes predisposes to development of cavernomas (Zhou *et al*, 2016). Somatic mutation of *MAP3K3* is also implicated in the molecular etiology of sporadic cerebral cavernomas (Weng *et al*, 2021). Normally, the CCM2/CCM3 complex binds and inactivates MAP3K3, but in a *Ccm2-/-* mouse model, vascular malformation can be prevented by genetic loss of *Map3k3* (Zhou *et al*., 2016). Notably, MAP3K3 activates both Rho (Zhou *et al*., 2016) and Hippo signaling (Lu *et al*., 2021), with the latter resulting in cytoplasmic retention of YAP/TAZ. It appears that the ALK1-SMAD4 and CCM axes may converge on similar pathways that interact with Hippo-YAP/TAZ and Cdc42 rho kinase activity. Of translational relevance, the use of MAP3K3 (Choi *et al*, 2018) and or Rho kinase inhibitors (Shenkar *et al*, 2019; Shenkar *et al*, 2017) may merit investigation for treatment of HHT.

Our gene expression network analysis highlights the PI3K pathway as associated with *Eng, Acvrl,* and *Ptpn14*. Notably, PI3K inhibitors are already in clinical trial for HHT (Alsina-Sanchís *et al*, 2018; Ola *et al*., 2018). Thalidomide and pomalidomide, drugs that target components of the ubiquitination machinery, are in clinical trial for HHT therapy (Snodgrass *et al*, 2021), and this class of IMiD drugs may affect both PTPN14/SMAD and PTPN14/YAP/TAZ ubiquitination and turnover. Future design of more specific drugs with fewer side effects, such as by targeting specific ubiquitin modifying enzymes, would benefit from knowledge of the interactions that regulate stability of PTPN14, SMAD4 and YAP/TAZ, and a deeper understanding of the physical interactions between signaling components of BMP9-ALK1-SMAD4, Hippo-YAP/TAZ, MAP3K3, and Rho GTPase Cdc42 pathways.

## Materials and Methods

### *Mus Spretus* × *Mus Musculus* F1 backcross and extraction of lung RNA

All animal experiments were approved *a priori* by the UCSF IACUC. 69 F1 backcross mice were generated by crossing inbred male SPRET/Ei with inbred female FVB/N mice (Jackson Laboratory). female F1 hybrids were then mated to male FVB/N mice to generate the backcross. Note that the male F1 mice of this inter-species cross are infertile. Lungs from eight-week-old mice were snap-frozen, and RNA was isolated usingTRIzol (Invitrogen) according to the manufacturer’s instructions. Residual contaminating genomic DNA was removed by DNase treatment (Ambion). RNA was extracted with Trizol followed by DNAse I treatment and purification (RNeasy RNA purification kit,Qiagen).The purified RNA was quantified with Nanodrop and QC checked by Agilent Bioanalyzer.

### Illumina Gene expression analysis

Gene expression was measured with the Illumina Mouse Whole Genome array (Illumina Inc.,San Diego,CA,USA). Sample preparations, hybridization and scanning were performed according to manufacturer’s instructions. Briefly, 200ng of total RNA was prepared to make cRNA by using Illumina Total Prep RNA Amplification Kit (Ambion). First-strand complementary DNA was generated with T7 oligo(dT) primer and Array Script and the second strand cDNA was synthesized using DNA polymerase. Biotin-NTP with T7 enzyme mixes was used to make biotinylated cRNAs. The labeled cRNAs were purified, quantified and checked again using the Bioanalyzer. The labeled cRNA target was used for hybridization and scanning according to Illumina protocols. Probes with present/absent call ≥0.05 assessed by Illumina Bead studio software, were marked absent. Raw microarray data were quantile normalized and log2-transformed. Statistical analysis was performed with Rversion2.13 (Team, 2020).

### Lung Gene Expression Correlation Analysis

Lung gene expression correlation analysis was undertaken using the program Carmen (Quigley *et al*., 2016).

### GO Enrichment Analysis

In order to test for functional enrichment in gene clusters, we implemented R/topGO v2.44.0 (Alexa, 2021) on Biological Process using Fisher exact tests, correcting for multiple testing with the Benjamini-Hochberg procedure, using annotations provided by R/org.Mm.eg.db v3.8.2 (Carlson, 2019). To capture an unbiased view of GO enrichment, in Figure 1D we display the top 10 most significantly enriched terms for each gene cluster according to FDR. We calculated enrichment scores as observed / expected within functional categories. We then centered and scaled across the matrix, generating a heatmap of the scaled scores using R/gplots/heatmap.2 v3.1.1 (Warnes *et al*, 2020).

### Cell culture

Human Umbilical Arterial ECs (HUAEC) from LONZA^®^ or PromoCell^®^ were used between passage 1 to 3. They were cultured in EC specific medium with 2% FBS (EGMTM-2 EC Growth Medium-2 BulletKit^TM^, LONZA^®^ or EC Growth Medium 2, PromoCell^®^) at 10^5^ cell/well in a 6 well plate. For over-expression studies, HEK293 cells (ATCC; CRL-1573™) were cultured in DMEM medium containing D-Glucose, L-Glutamine and Sodium pyruvate (GIBCO^®^), 10% FBS and 1% Penicillin/Streptavidin. TMLC cells are a stable subclone of CCL64 mink lung epithelia cells, stably with an expression construct containing a TGF-b-responsive promoter fragment, consisting of a truncated plasminogen activator inhibitor-1 (PAI-1) promoter, fused to the firefly luciferase reporter gene (Abe *et al*, 1994). MG132 (M7449, SIGMA-Aldrich^®^) at 1µM was used in the culture media for proteosome inhibition, and cycloheximide at 10ug/ml was used for inhibition of protein translation. In each case, cells were treated with drug for 24 hours before the protein extraction.

### Gene Expression Knock Down (KD)

When the cells were 80% confluent, they were transfected with 0.006 pMol of *siRNA* (Dharmacon®) for each gene under study using RNAi Max Lipofectamine (Invitrogen®) (**Supplementary Table S5**). 48h after transfection, cells were treated with 2 ng/ml BMP9 (R&D®) with no prior serum starvation since serum starvation reduced SMAD4 levels in HUAEC. At the stated end points post-treatment, cells were washed with PBS and lysed (200 µl) with M-PER Mammalian Protein (ThermoScientific®) plus 1x Protease inhibitor (cOmplete Mini EDTA, Roche®). Cells were detached using a scraper (on ice), sonicated to lyse for 3s (20 kHz, using the Q500 Sonicator). After centrifugation of 13, 000 rpm for 10 min at 4°C, the supernatant was collected and stored at −20C for analysis.

### Growth factor signaling

Confluent HUAEC cells in EC specific medium with 2% FBS were treated with 2 ng/ml BMP9 (Recombinant Human BMP-9 Protein, # 3209-BP, R&D®) and proteins extracted at specified time points after treatment. BMP9 signaling inhibition was performed using an ALK1 kinase small molecule inhibitor K02288 5 µM (#S7359, Selleckchem®). TGFβ treatment was undertaken using 2ng/ml of Recombinant Human TGF-beta 1 Protein (#240-B, R&D®) and signaling specificity tested using the specific inhibitor for ALK5 kinase activity: SB431542 5 µM (#S1067, Selleckchem®).

### Ectopic protein expression

HEK293 cells were cultured in DMEM (GIBCO®) with 10% FBS (VWR®) and 1% Penicillin/Streptavidin. The V5-tagged PTPN14 expression constructs described by Liu et al. (Liu *et al*., 2013) were used for PTPN14 over-expression studies. Flag-tagged SMAD4 (#80888, Addgene®), and HA tagged TAZ (#32839, Addgene®) cloned into pcDNA, were also used. Cells were transfected with 2.5 µg of plasmid per well of a 6-well plate using Lipofectamine 3000 (Invitrogen®) and protein expression levels analyzed 24h after transfection.

### Western Blot

Western blot analysis was undertaken using 7.5% TGX gels (BioRad®) transferred onto 0.45 µm PVDF membranes (Amersham™ Hybond™, GE life science®). Transfer was undertaken using the BioRad® semi-dry system. After blocking the membrane for 1h at RT with AdvanBlock (Advansta®), and proteins were detected by incubation with relevant antibodies (**Supplementary Table S6**) diluted 1/500 in 2% BSA, followed by HRP-tagged secondary antibody (1/5000), in TBST 1x (CellSignaling®). Signal was detected using SuperSignal™ West Femto Maximum Sensitivity Substrate (Thermo Scientific™), and images captured using a ChemiDoc Imaging System (BioRad^®^).

### Co-immuno-precipitation

Protein (10 µg) was immuno-precipitated using a target antibody (1 µg) with Pierce™ Protein A/G Magnetic Beads. Protein elution was undertaken using 1x LD Laemmli Sample Buffer (BioRad^®^) and denaturized at 95°C for 5 minutes before transferring to ice before gel loading.

### Nuclear cytoplasmic fraction

Cell fractionation was performed using NE-PER™ Nuclear and Cytoplasmic Extraction Reagents (Thermo Fisher™) and lysates were analyzed by Western-blot and/or used for Co-IP.

### Dual Luciferase activity test

HEK293 cell were transfected with pGL3BRE Luciferase (#45126, Addgene^®^) and pCMV-Renilla luciferase plasmid (Thermofisher #16153). Cells were transiently transfected with PTPN14 V5 tagged expression constructs (Liu *et al*., 2013), after 24 hours cell are treated with BMP9 (2ng/ml) or combined to K02288 at 1µM (ALK1 kinase inhibitor and relative luciferase activities quantified after 16 hours using the Dual-Glo^®^ Luciferase Assay System (Promega^®^). The TGFβ Reporter Assay was performed in TMLCs after transfection of the various PTPN14 constructs, and luciferase activity was measured following 16h treatment with TGFβ (2ng/ml) or SB431542 (1µM) (ALK5 kinase inhibitor) and relative luciferase activities quantified after 16 hours using the Dual-Glo^®^ Luciferase Assay System (Promega^®^). Histograms were generated and data analyzed by multiple paired t-tests and 2-way ANOVA tests using GraphPad (PRISM^®^ 9.0.0).

### Proximity Ligation Assay (PLA)

PLA was carried out on confluent, low passage primary HUAEC monolayers using the Duolink Proximity Ligation Assay (PLA) from Sigma. Cells were treated with 2 ng/mL BMP9 in cell culture media for one hour at 37°C. Controls remained untreated. Cells were then washed with PBS, fixed with 4% paraformaldehyde in PBS for ten minutes, and permeabilized in 0.2% Triton X-100 for ten minutes at room temperature. Cells were blocked in PLA blocking buffer for one hour at 37°C. Anti-PTPN14 (Sigma Prestige), anti-pSMAD1/5 (Cell Signaling), anti-SMAD4 (Invitrogen) antibodies were used as primaries. Cells were counterstained with DAPI for nuclear stain. Phalloidin was used to stain cell membranes. Imaging at 40X was done using a Keyence microscope. Cytoplasmic and nuclear PLA signals per cell were quantified using CellProfiler^TM^ software. Three separate repetitions were grouped for analysis. Histograms were generated and data analyzed for statistical significance by multiple paired *t*-test using GraphPad (PRISM 9.1.0).

### RNAScope *in situ* hybridization

*In situ* hybridization was undertaking using standard protocols and reagents from the RNAScope 2.5 HD Duplex Detection Kit™ (Advanced Cell Diagnostics). Wild type C57BL/6J 8.5 dpc embryos, and adult lung and heart were fixed in 4% paraformaldehyde and embedded in paraffin. 5uM sections were used. Differentially labelled RNA-specific oligonucleotide probes for *Eng* and *Ptpn14* were hybridized simultaneously, and chromogenic signals were detected at 10X and 40X by standard bright-field microscopy using a Keyence microscope.

### Immunofluorescent staining

Lung tissue was harvested and fixed at 4°C in 4% paraformaldehyde in PBS before being dehydrated in ethanol and embedded in paraffin. 5uM sections were used. Hematoxylin and eosin staining was performed per standard protocol. For fluorescent staining, the tissue was cleared with xylene, dehydrated with ethanol, permeabilized with .1% Triton-X 100, and blocked with 1% BSA, 20% goat serum in PBS overnight at 4°C. Anti-Ptpn14 (Sigma Prestige) and anti-CD31 (BD Pharmingen) antibodies were applied concurrently in blocking solution overnight at 4°C. Tissues were washed with PBS and incubated in Alexa Fluoro 488 and 647 (Invitrogen) concurrently in blocking solution for two hours at room temperature in a darkened chamber. Tissues were counterstained with DAPI for nuclear stain and slides were mounted with ProLong Gold with DAPI (Invitrogen). Tissue imaging at 10X and 40X was carried out using a Keyence microscope.

## Acknowledgments: Sources of Funding, & Disclosures

We thank Rik Derynck for critical pre-review of the manuscript, and Adila Izgutdina for laboratory assistance. The work was funded by NIH grants R01-HL122869 and R01CA210561. OM was funded in part by a fellowship from the ADDAX foundation. DTB was funded in part by a University of California, Presidential Post-doctoral Fellowship.

## Authors contributions

OM, DTB, RJA, MT, designed aspects of the project, interpreted the data, and wrote the manuscript. AB and MT generated the mouse-specific backcross and lung microarray expression data. OM, MT, AB and RJA undertook bioinformatics analysis. SB, XC, and ST provided technical support. JZ provided PTPN14 expression constructs. AB and JZ contributed to wirting the manuscript.

## Disclosure and competing interests’ statement

The authors have no financial disclosures relevant to this study.

## SUPPLEMENTARY FIGURE LEGENDS

**Supplementary Figure S1: *PTPN14* is preferentially expressed in ECs of human and mouse lung.**

Single cell RNAseq data from LungMap (https://lungmap.net/) was interrogated for *PTPN14* expression. **A**: Relative *PTPN14* RNA levels in sorted human lung cells, specified by cell type, and at the indicated ages post-partum. Green, endothelial cells; yellow, epithelial cells; orange, mesenchymal cells; blue, immune cells. **B**: Relative *Ptpn14* expression levels in sorted cells of mouse lung 28 days post-partum, with highest levels observed in ECs > fibroblasts > B lymphocytes and undetectable levels in macrophages and epithelial cells.

**Supplementary Figure S2. Negative controls for chromogenic RNAScope *in situ* hybridization.** Negative control for RNAScope™ using non-specific RNA probes on (**A**) lung or **(B)** heart, at 10X (left) or 40X (right). All tissues were counterstained with hematoxylin.

**Supplementary Figure S3: *PTPN14* KD reduces SMAD4 and endoglin protein levels. A:** Western blot analysis of HUAEC protein lysates analyzed at different cell densities to show increased PTPN14 and endoglin expression level as cells attain higher confluence. **B:** Western blot of protein lysates from HUAECs 48 hours after transfection with two siRNAs against *PTPN14* and two against *SMAD4,* compared to non-targeted siRNA. This confirms the destabilizing effect of *PTPN14* KD on SMAD4 and endoglin protein levels. **C:** Western blot analysis of HUAEC protein lysates transfected with *siNT, siSMAD4* or *siPTPN14*, followed by treatment for 0, 1 or 24h with 2 ng/ml of BMP9. *siPTPN14* reduces expression of both SMAD4 and endoglin. Consistent with the *ENG* gene being a transcriptional target of SMADs, *siSMAD4* obliterates endoglin expression. In contrast it enhances expression of PTPN14 in a BMP9 dependent, SMAD4-independent manner.

**Supplementary Figure S4: In HEK293 cells, PTPN14 binds and stabilizes SMAD4 through the PTPN14 linker region, whereas it reduces TAZ protein levels. A:** Western blot of HEK293 protein lysates shows high ALK1 expression levels, that are reduced by *siALK1*, demonstrating specificity of western. **B**: HEK293 cells were transfected with HA-tagged TAZ, with or without knockdown of endogenous *PTPN14* using *siPTPN14*. Western blot analysis shows that TAZ-HA decreases PTPN14 levels, whereas *siPTPN14* elevates expression of ectopic TAZ-HA, as detected using an anti-TAZ antibody. **C**: Western blot analysis of nuclear and cytoplasmic protein lysates of HEK293 cells transfected with *siPTPN14*, V5-PTPN14^-wt^, or V5-PTPN14^-ΔFERM^ combined with TAZ-HA, shows that ectopic PTPN14 expression has little effect on nuclear levels of TAZ-HA, but *siPTPN14* enhances cytoplasmic TAZ-HA levels. V5-PTPN14-WT decreases TAZ-HA in a dose dependent manner in the cytoplasm but not in the nucleus. Co-transfection of PTPN14^-ΔFERM^ has a similar effect as PTPN14-WT, showing that this mutant is not defective in turning over TAZ. **D:** Western blot analysis of HEK293 cell lysates after ectopic expression of the various WT and mutant V5-PTPN14 expression constructs with SMAD4 to further demonstrate stabilization of SMAD4 by PTPN14, which is most effective for the PTPN14-PPXY mutant construct. This lysate was used for reciprocal co-immunoprecipitation using anti-V5 or anti-FLAG antibodies. All constructs showed binding between PTPN14 and SMAD4, suggesting that SMAD4 binds to the PTPN14 linker region, independently of the two PPXY-binding sites that are absent in the PPXY-PTPN14 mutant.

---

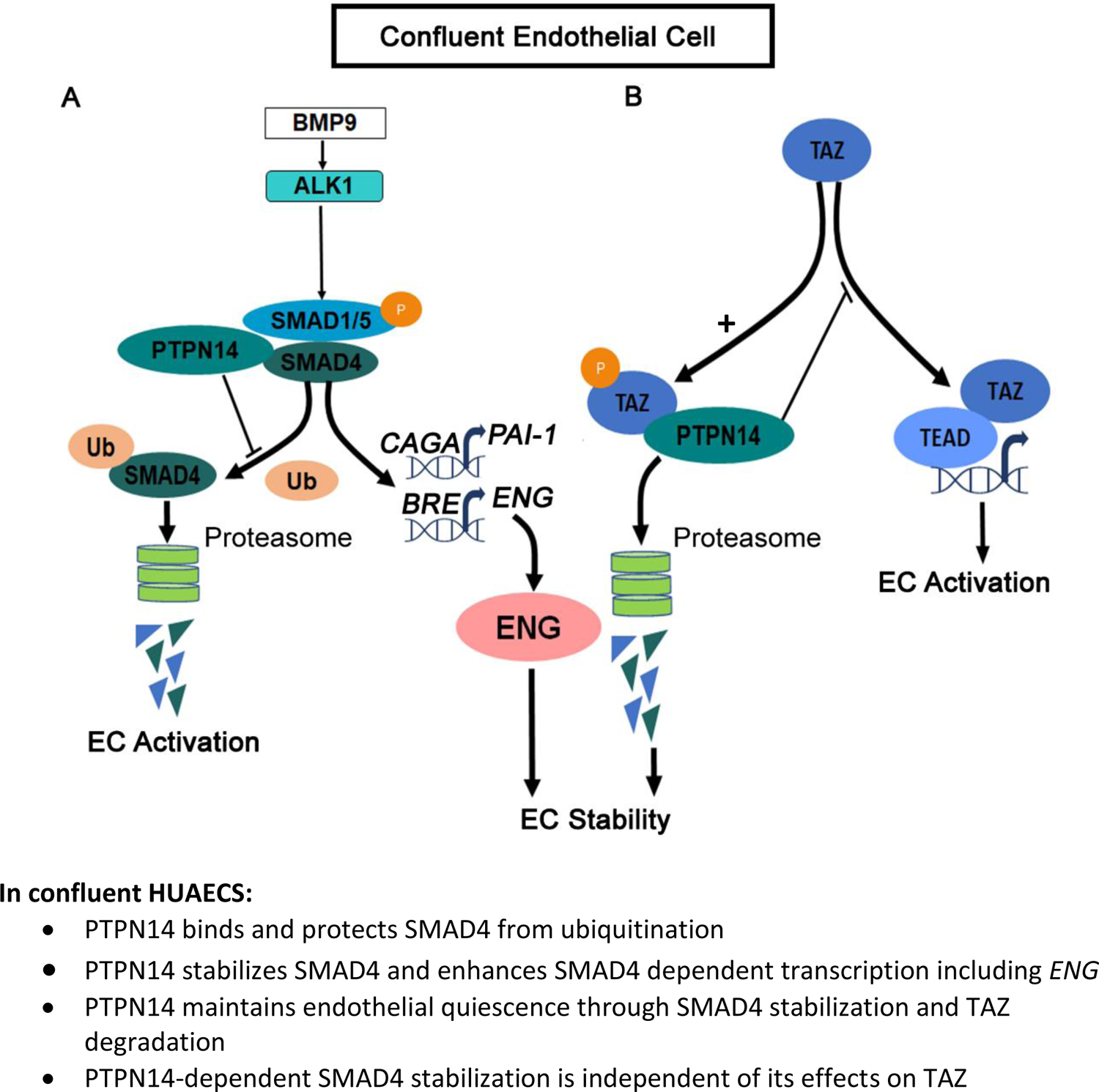

## Notes

### Competing Interest Statement

The authors have declared no competing interest.

