## Supplementary figures and images for "PTPN14, a modifier of HHT, protects SMAD4 from ubiquitination and turnover to potentiate BMP9 signaling in endothelial cells"

### Supplementary Figure S1

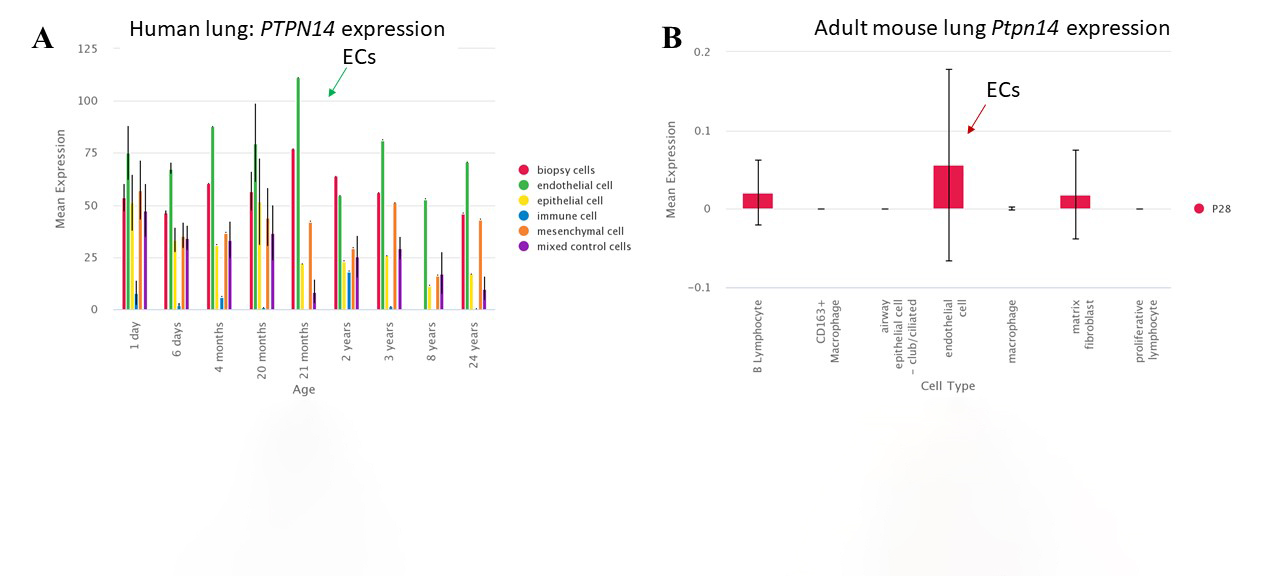

### Supplementary Figure S2

A

B

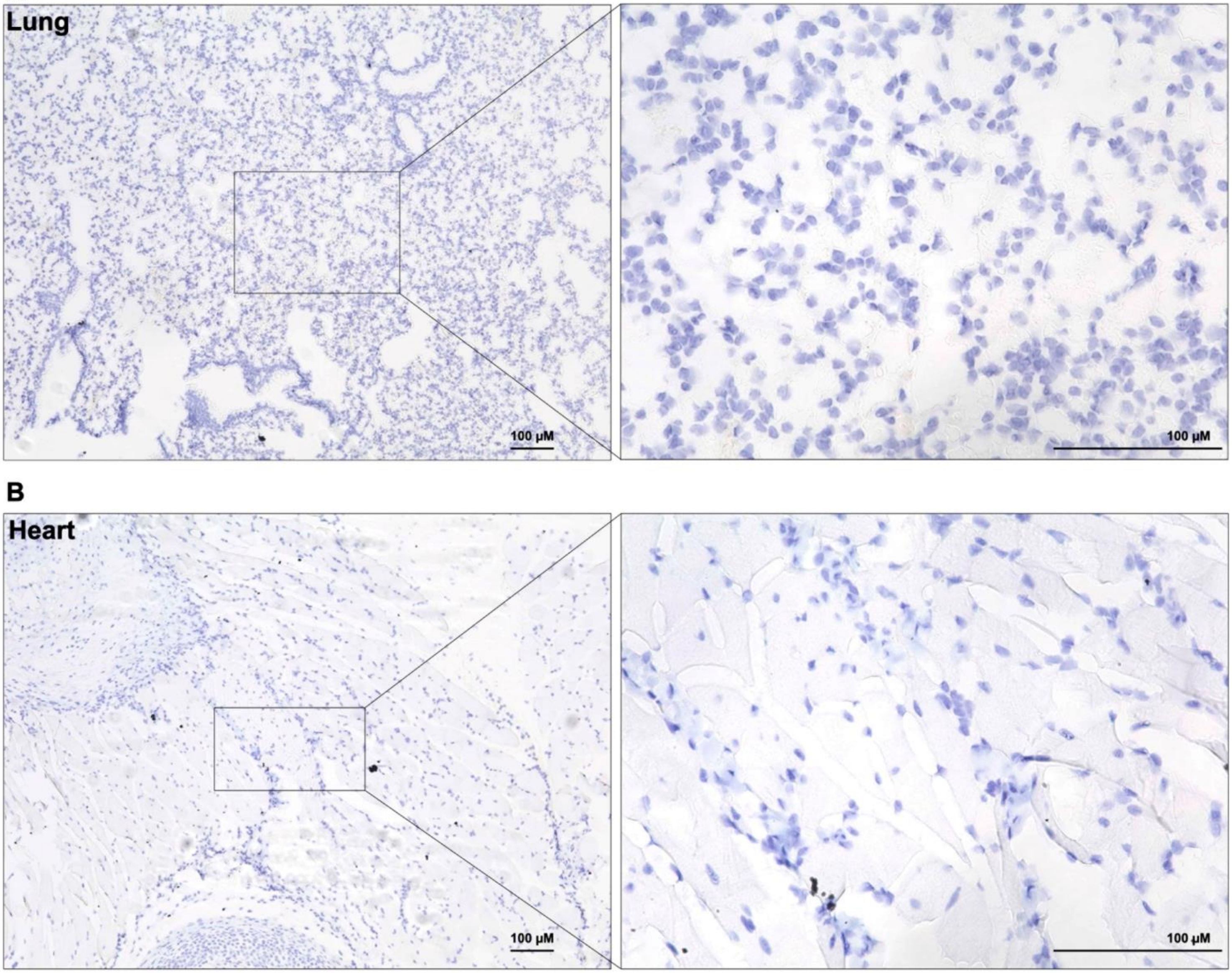

### Supplementary Figure S3

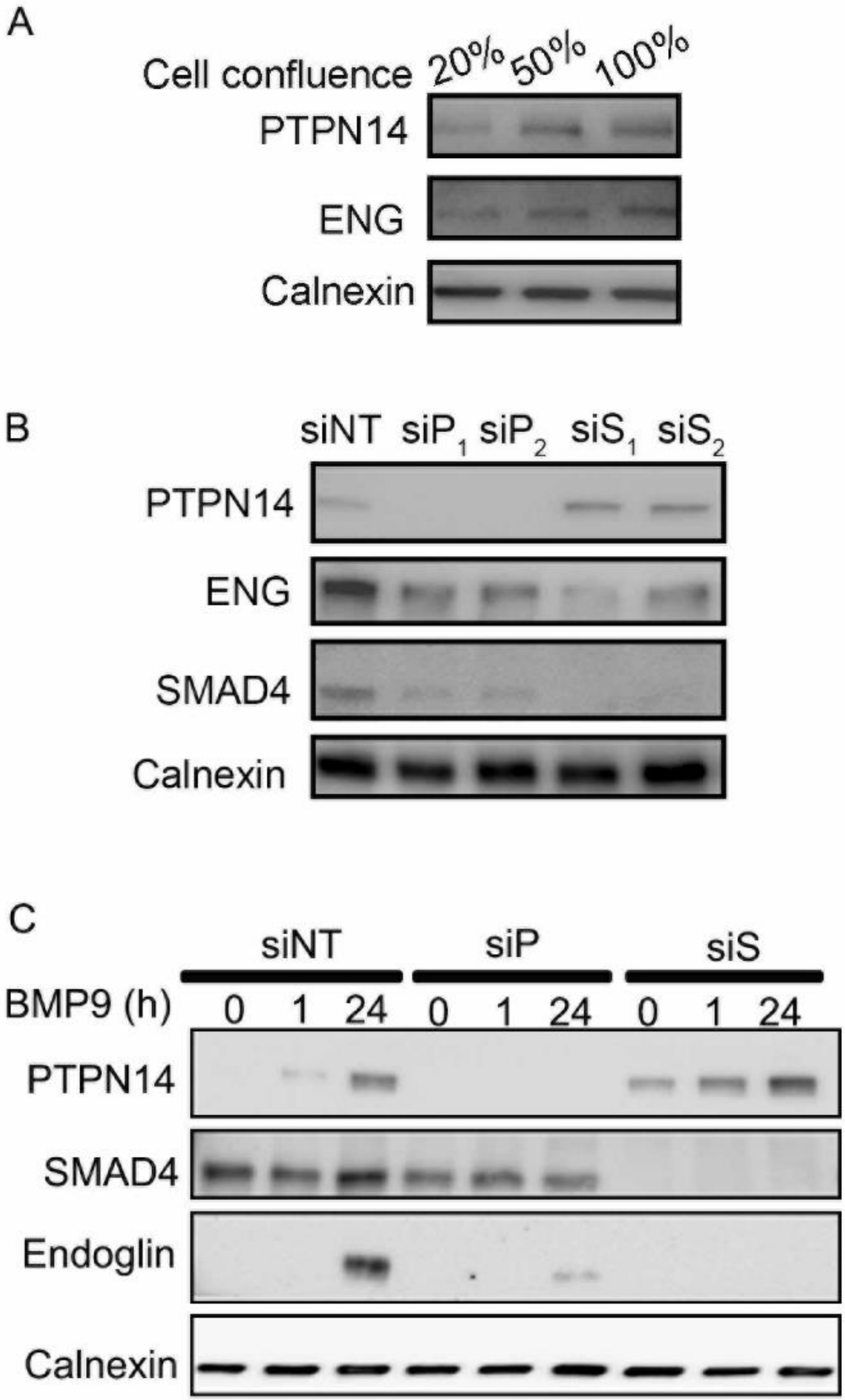

### Supplementary Figure S4

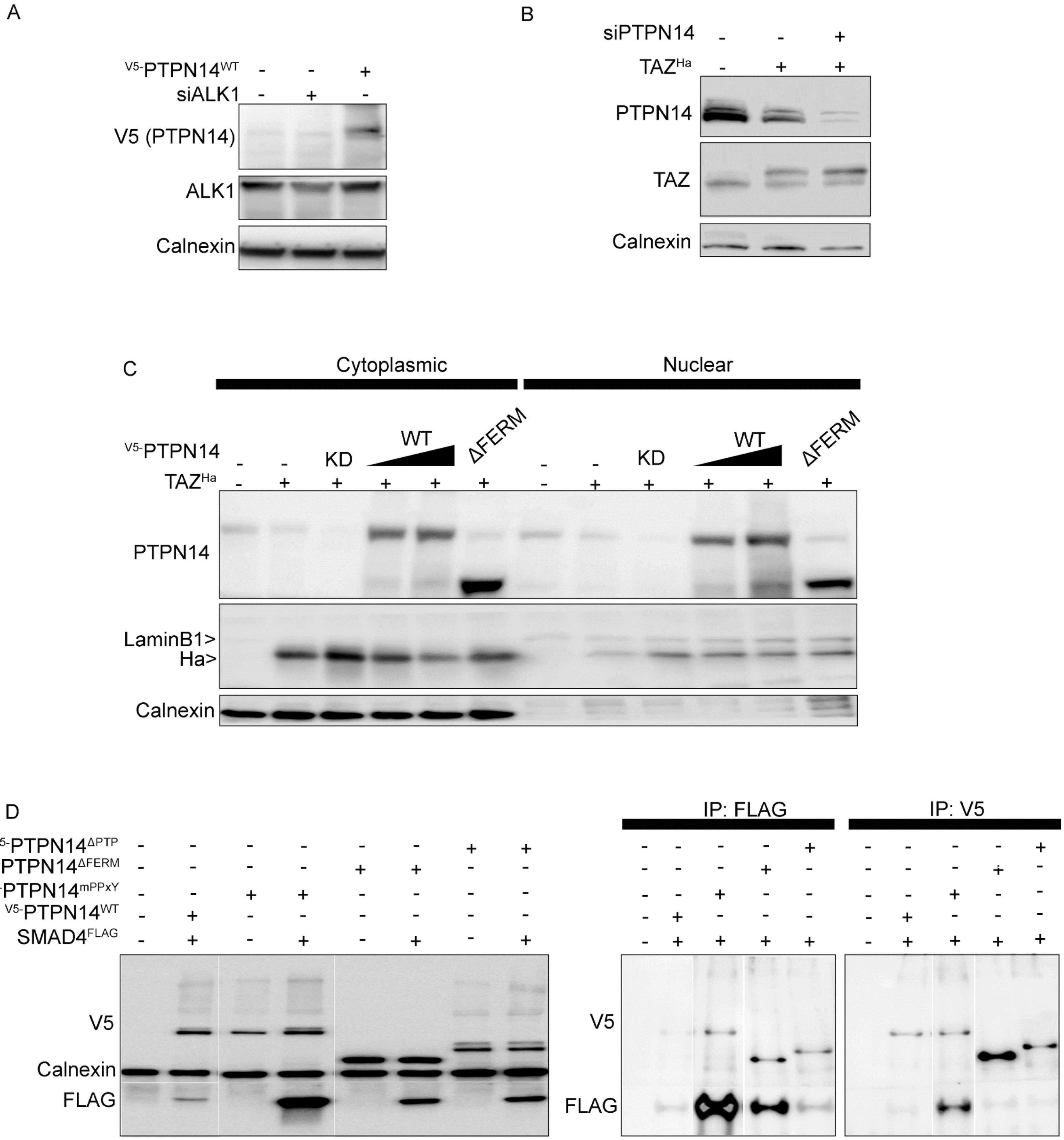
